# The immune landscape of pediatric Brain and Solid Tumors: A systematic review and integration of immunohistochemistry and single-cell RNA sequencing data

**DOI:** 10.1101/2025.06.06.658323

**Authors:** Francisca J Bergsma, Jan Koster, Bob Baalman, Francis Mussai, Hubert N Caron, Jan J Molenaar, Judith Wienke

## Abstract

**Background:** Immunotherapies achieved remarkable success in adult cancers, yet their efficacy in pediatric brain and solid tumors remains limited. Insights into the unique immune landscape of pediatric tumors are crucial to improve immunotherapies for pediatric patients.

**Methods:** We performed a systematic search for studies reporting immunohistochemistry (IHC), single-cell- or single-nucleus RNA-sequencing (sc/snSeq) data on the immune landscape of pediatric tumors. For IHC studies, data on macrophages, T cells, T helper cells, regulatory T cells, NK cells and B cells were extracted. For sc/snSeq studies, cell cluster counts were extracted. 47 IHC studies and 26 sc/snSeq studies were included in the analysis.

**Results:** Our integrative in-depth analysis of 73 studies covered 35 unique pediatric tumor types with 17 tumor types analyzed by IHC, 4 by sc/snSeq, and 14 by both techniques. Regardless of variability in analysis methods, both IHC and sc/snSeq showed that peripheral nerve tumors and soft tissue sarcomas had relatively immune-infiltrated, T cell-rich tumor microenvironments (TME). Brain tumors exhibited a macrophage/microglia-rich, NK cell-infiltrated and T cell-depleted TME. Sc/snSeq data confirmed these observations, showing a macrophage/microglia-rich brain TME. Compared to adult tumors, (CD8^+^) T cell infiltration and macrophage infiltration was low for all pediatric tumor types. Integrated IHC and sc/snSeq data were visualized in interactive heatmaps, publicly available on R2 as a comprehensive atlas [https://hgserver1.amc.nl/cgi-bin/r2/main.cgi?option=imi2_targetmap_v1; map Immune_landscape_mm2(by patient)_v4].

**Conclusion:** We provide a comprehensive, integrated overview of the immune landscape of pediatric brain and solid tumors. These insights can aid the development and selection of immunotherapeutic strategies for specific pediatric cancers, tailored to their unique immune characteristics.

## Introduction

Pediatric cancers are the leading cause of death among children and adolescents under the age of 20 [1]. Solid and brain tumors account for 65% of all pediatric cancers and they encompass tumor types with the poorest prognosis like high grade glioma, ependymoma, rhabdoid tumors, neuroblastoma, and osteosarcoma [1–3]. Advances in diagnostic techniques and cancer therapy such as radiotherapy and chemotherapy, have significantly improved the average 5-year survival rate of pediatric patients with solid cancers up to 80% [3–5]. However, challenges remain in achieving optimal long-term survival and minimizing therapy induced toxicity, as childhood cancer survivors are at high-risk of developing late complications including heart disease, lung problems, secondary malignancy, and infertility, which significantly impact their quality of life [6].

Immunotherapy has emerged as a promising therapeutic strategy for cancers by redirecting the body’s own immune system to recognize tumor cells and induce tumor killing. Nevertheless, as compared to adult cancers, the success of immunotherapies in pediatric tumors is limited and varies between different tumor types. Especially in solid and brain tumors the development and success is very limited [5,7,8]. To date, only three immunotherapeutics have been approved and marketed for pediatric solid tumors, namely Dinutuximab, a monoclonal antibody for high-risk neuroblastoma patients, Ipilimumab, a checkpoint inhibitor for patients with advanced melanoma and Pembrolizumab, a checkpoint inhibitor for a variety of pediatric solid tumors [9]. The main challenges for the development of novel immunotherapeutic strategies for pediatric solid and brain tumors are the relatively low tumor mutational burden (TMB) as compared to adult tumor, low MHC-I expression in some tumor types, and the highly immunosuppressive tumor microenvironment (TME) [7,10]. The low TMB results in a low expression of neoepitopes, and low MHC-I expression on the tumor cell surface further limits the recognition by T cells [8],[9]. The limited T cell infiltration reduces the efficacy of immunotherapies that mainly rely on T cell presence, such as bispecific T-cell engagers (BiTEs) [13], and immune checkpoint blockade (ICB) [11,14]. The pediatric solid TME is characterized by a high presence of immunosuppressive tumor-associated macrophages (TAMs) [15], which are known to reduce the efficacy of chimeric antigen receptor (CAR) T cell therapies [16]. Thus, while T cell-based immunotherapies face challenges in pediatric solid and brain tumors due to their low immunogenicity and highly immunosuppressive TME [11], alternative approaches that rely on macrophages or NK cells, such as antibody-dependent cellular phagocytosis or cytotoxicity (ADCP/ADCC)-based therapies may offer potential benefits [17,18]. Given the highly heterogenous immune landscapes across pediatric tumors, immunotherapeutic strategies should be selected based on the specific characteristics of the TME to enhance the efficacy.

Understanding the unique TME composition of pediatric brain and solid tumors is crucial for enhancing the efficacy of immunotherapies and tailoring treatment strategies to specific tumor types. Here we present a systematic review of the immune landscape in pediatric solid and brain tumors by integrating data from immunohistochemistry (IHC) and single-cell/single-nucleus RNA-sequencing data (sc/nSeq). This comprehensive overview may aid future tailoring of immunotherapies for specific pediatric cancers, with a focus on optimizing immunotherapeutic treatment strategies by taking full advantage of the unique characteristics of the pediatric TME.

## Methods

### Search strategy

The systematic review was conducted following the Preferred Reporting Items for Systematic Reviews and Meta-Analyses guidelines [19]. IHC and sc/snSeq studies were identified by a computerized search through the PubMed search database with no date restriction up to November 13, 2024.

Two separate searches were conducted: one for the IHC data and another for sc/snSeq data. Both search strategies were based on three main components: technique, pediatric focus and cancer subtypes. To refine the IHC search for immune cells, the term (“immune “[All fields]) was included. A more detailed description of the full search strategy is included in **Supplementary Table 1**. To enable comparison with pediatric tumors, selected adult cancer studies were included, identified through targeted but non-systematic searches.

### Study selection

Before screening, duplicated studies and non-English reports were removed. Eligible studies were included if the following criteria were met: (1) the cohort had a mean age or age range below 25 years, (2) tumor subtypes were analyzed separately, (3) human tumor samples were used, (4) sc/snSeq data were newly generated by the study, (5) data were analyzed and either reported in the study or provided upon request.

Studies were excluded if they were reviews, meta-analyses, case reports, protocols, preprints, abstracts, previews, chapters or corrections.

### Data extraction

The following descriptive variables were extracted from eligible studies: first author, year of publication, PMID, tumour type, cohort size, mean age or age range, samples origin, tissues condition and treatment regime.

For IHC-specific studies, additional methodological details were collected, including the scoring method, pathologically confirmed tumour tissue used, use of proper antibody controls, staining approach (manual or automated, single or multiplex), and image assessment method (manual or digital). These details were used to calculate an overall quality score for each study, based on six predefined criteria, with each criterion weighted by its own importance factor (**Supplementary Table 2 and 3**). The resulting score, ranging from 0 to 15, serves as an indicator of the robustness and validity of the reported findings.

For the sc/snSeq dataset, we recorded whether the study used single-cell or single-nucleus RNA sequencing and whether samples were enriched for specific cell types.

### Single-cell and single-nucleus RNA sequencing cell subtypes

In order to facilitate comparisons between different pediatric tumors, we grouped specific cellular subtypes into overarching groups (**Supplementary Table 4**). As total cell count we used the total number of cells reported, with omission of red blood cells and erythrocytes. The category “other cells” was defined as the sum of glial cells, schwann cells/stroma, astrocytes, oligodendrocytes, ependymal cells, neuronal cells, pericytes, stellate cells, progenitors cells and other cells. “Immune cells” where defined as the sum of lymphoid, myeloid and hematopoietic stem and progenitor cells (HSPCs). “Lymphoid cells” were defined as the sum of T cells, T/NK cells, NK cells, ILCs, B cells and plasma cells. “Myeloid cells” were defined as the sum of mast cells, macrophages, monocytes, neutrophils, mDCs and p/cDCs.

### Quantification and statistical analysis

In order to enable fair comparisons of marker expression or cell type numbers between studies with similar or identical pediatric tumor types, but differing cohort sizes, we calculated the weighted mean expression for each marker per tumor type based on the cohort size: *weighted mean = (mean^study1^ x cohort size^study1^ + mean^study2^ x cohort size^study2^)/cohort size^study1+study2^*.

Principal Component Analysis (PCA) was performed on scRNAseq derived proportions of tumor, mesenchymal and fibroblasts, endothelial, immune, and other cells using the *PCA()* function from the FactoMineR package in R (version 4.4.0). Visualization was generated using the *fviz_pca_biplot()* function from the factoextra package.

All statistical analyses were conducted in R (version 4.4.0). Differences in immune cell densities between tumor categories were assessed using the Kruskal-Wallis test, followed by Dunnett’s post-hoc test for pairwise comparison with Bonferroni P-value correction for multiple testing. Linear regression analysis was performed using the *lm* function from the ‘stats’ R package to evaluate correlations between immune subsets. Statistical significance was defined as p < 0.05.

## Results

### 1. Characteristics of the included studies

The systematic literature search in PubMed identified 722 studies (306 for IHC and 416 for sc/snSeq), of which 612 studies (275 for IHC and 337 for sc/snSeq) underwent full review (**Fig. 1a and b**). After careful screening, 47 IHC studies and 26 sc/snSeq studies met all eligibility criteria and were included in our systematic review. Of note, 42 sc/snSeq studies which met all inclusion criteria, did not provide the number of cells per cluster and could therefore not be included. Furthermore, the scoring methods used in different IHC studies varied substantially, thereby hampering direct comparison between studies. To address this, we categorized the different scorings methods into 4 main methods: cells per mm^2^, cells per field, percentage positive cells per field, and percentage of positive samples. Scoring methods that did not fit into these categories were classified as “other”, including studies using H score [20], IHC score [21], low/high [22–24], and percentage of high cases [25]. The full characteristics of the included studies are detailed in **Supplementary Table 3 and 4**. Additionally, the concatenated cells per mm^2^ IHC data as well the sc/snSeq are publicly accessible via R2 (https://hgserver1.amc.nl/cgi-bin/r2/main.cgi?option=imi2_targetmap_v1; map Immune_landscape_mm2(by patient)_v4).

**Figure 1.**
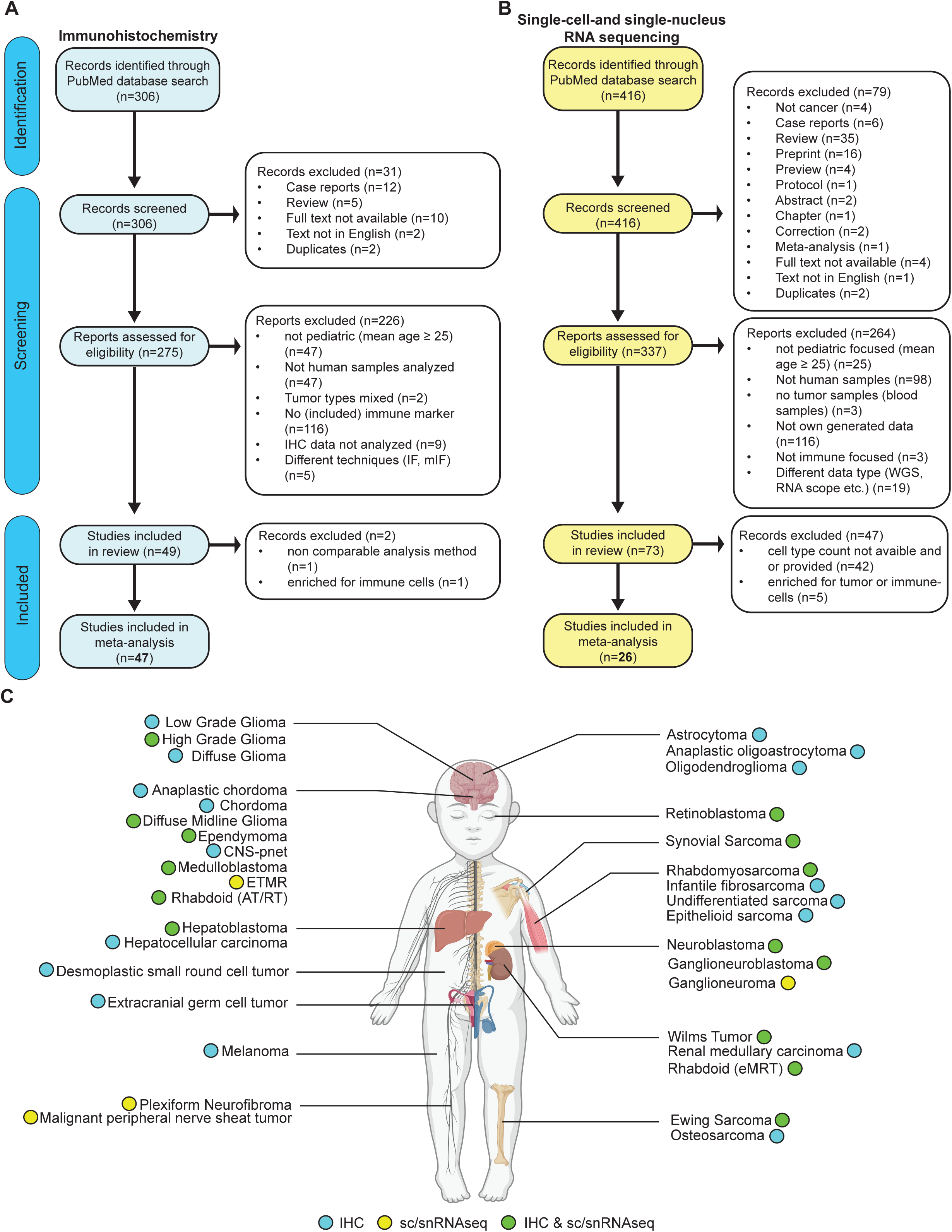
Pediatric brain and solid tumor cohort. (**a-b**) Preferred Reporting Items for systematic reviews for Systematic Reviews and Meta-Analyses guidelines of the selection of studies to be included for IHC (**a**), and sc/snSeq (**b**). (**c**) Overview of the 35 pediatric tumor types included in the systematic review. Figure made with Biorender.com.

In total, the included studies comprised 35 different pediatric tumor types, of which 17 were analyzed by IHC, 4 only by sc/snSeq, and 14 were covered by both techniques (**Fig. 1c**). We did not include the “other” IHC analysis category in the integrated analysis in this manuscript.

### 2. Lymphoid landscape across pediatric brain and solid tumors assessed by IHC

We first examined the presence of lymphocytes across pediatric tumors, focusing specifically on T cells (CD3^+^), B cells (CD20^+^), and NK cells (CD56^+^, CD57^+^, NKp46^+^, and NCR1^+^).

#### 2.1. T Cell-Rich Tumor Microenvironment In Soft Tissue and Peripheral Nerve Tumors

Across the 47 included studies, 12 studies analyzed lymphoid infiltration using a quantitative scoring method, expressed as positive cells per mm^2^ [26–37]. Overall, CD3^+^ T cells were present in the TME of most tumor types, with a median of 48 cells/mm^2^ (range: 0-398 cells/mm^2^). The intermediately malignant tumor ganglioneuroblastoma showed the highest T cell density compared to more malignant tumors (**Fig. 2a**). In rhabdomyosarcoma we found the second highest T cell infiltration, though at ∼50% of the level observed in ganglioneuroblastoma. Considerable variability in T cell levels was observed across rhabdomyosarcoma studies, which could not be attributed to differences between the embryonal and alveolar subtypes (**Fig. 2b**). Rather, variations in the analysed tissue types (e.g. primary versus metastatic tumors) may have contributed to the observed differences. Neuroblastoma, exhibited T cell densities similar to rhabdomyosarcoma with limited variation across studies (range: 139-212 cells/mm^2^; **Fig. 2a**). Importantly, MYCN-amplified neuroblastoma tumors, associated with a poor prognosis, exhibited lower T cell infiltration than MYCN non-amplified tumors (63 vs 241 cells/mm^2^; **Fig. 2c**), aligning with previous findings [38,39].

**Figure 2.**
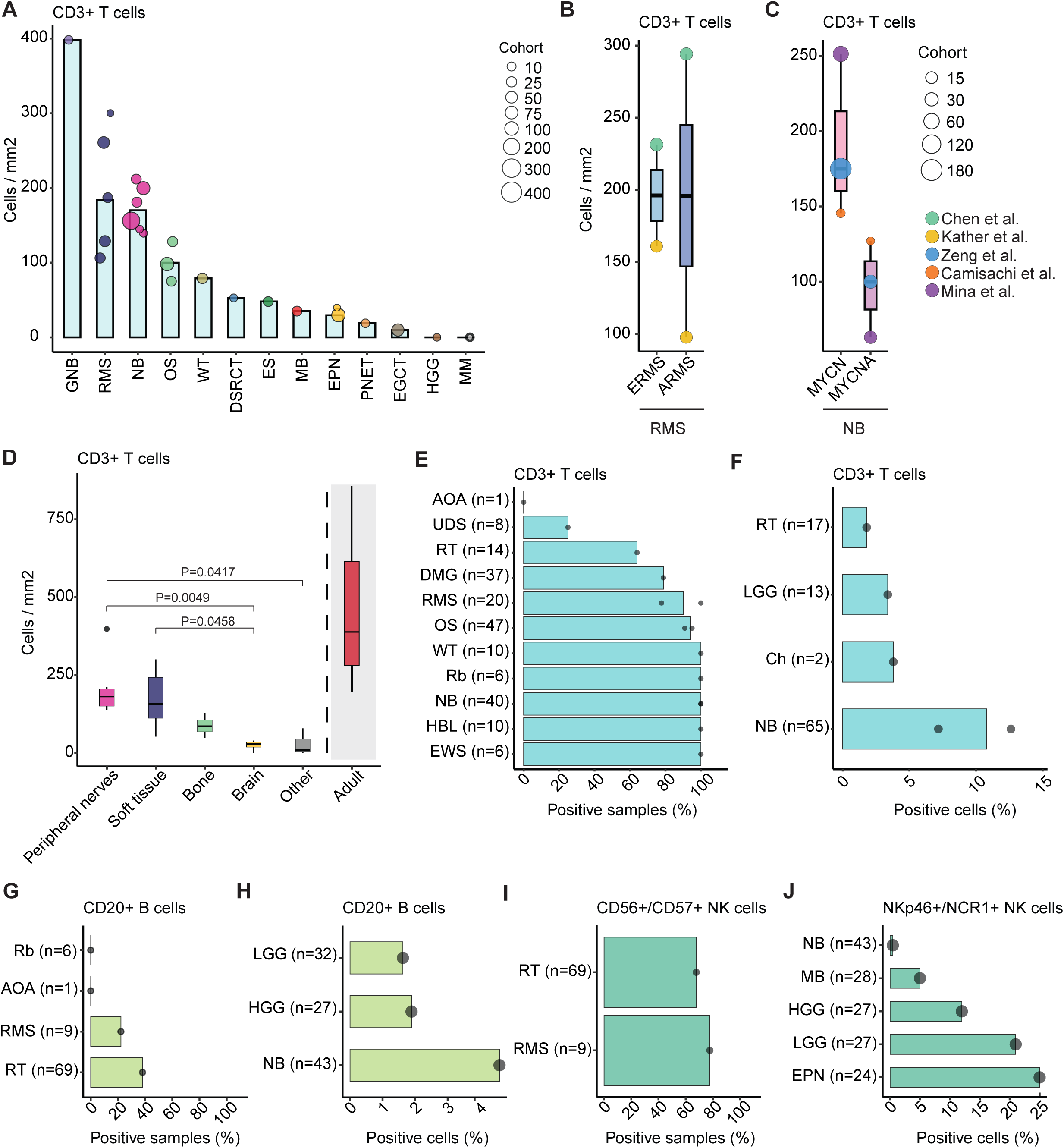
Lymphoid immune landscape across pediatric brain and solid tumors. (**a**) Bar plot of weighted mean CD3^+^ T cell infiltration per mm^2^ across pediatric (n=13) tumors, with individual (n=12) study distributions. (**b**) Median CD3^+^ T cell infiltration per mm^2^ in the embryonal (ERMS) and alveolar (ARMS) rhabdomyosarcoma (RMS) subtypes. (**c**) Median CD3^+^ T cell infiltration per mm^2^ in the MYCN non-amplified (MYCN) and MYCN-amplified (MYCNA) neuroblastoma (NB) subtypes. (**d**) Median CD3^+^ T cell infiltration per mm^2^ across tumor categories, based on individual studies. Statistic Kruskal Wallis with Dunn’s and Bonferroni P-value correction was used. (**e**) Percentage of CD3 positive tumor samples across pediatric (n=11) tumors. (**f**) Percentage of CD3 positive T cells across pediatric (n=4) tumors. (**g-h**) Percentage of CD20 positive tumor samples (**g**), and percentage of CD20 positive B cells (**h**) across pediatric tumors. (**i-j**) Percentage of CD56/CD57 positive tumor samples (**i**), and percentage of NKp46/NCR1 positive NK cells (**j**) across pediatric tumors. Abbreviations are as follows: MB, medulloblastoma; HGG, high-grade glioma; LGG, low-grade glioma; PNET, primitive neuroectodermal tumors; EPN, ependymoma; RMS, rhabdomyosarcoma; DSRCT, desmoplastic small round cell tumors; NB, neuroblastoma; GNB, ganglioneuroblastoma; OS, osteosarcoma; EWS, Ewing sarcoma; EGCT, extra-cranial germ-cell tumor; WT, Wilms tumor; MM, multiple myeloma; RT, Rhabdoid tumor; AOA, anaplastic oligoastrocytoma; Ch, chordoma; Rb, retinoblastoma; UDS, undifferentiated sarcoma; DMG, diffuse midline glioma; HBL, hepatoblastoma .

In contrast to the T cell-rich TME observed in rhabdomyosarcoma and neuroblastoma, brain tumors exhibited an overall lower T cell infiltration, ranging from 0-35 cells/mm^2^ (**Fig. 2a**). Moreover, when grouping the tumor types into larger tumor classes, brain tumors demonstrated significantly lower T cell densities compared to peripheral nerve and soft tissue tumors (**Fig. 2d**). Notably, even the highly infiltrated pediatric tumor types had a ∼2 fold lower T cell infiltration than adult tumors (range of adult tumors with both low and high TMB). This suggest that T cell infiltration is generally lower in pediatric tumors compared to adults, highlighting potential challenges in directly translating immunotherapies from adult to pediatric cancers.

In addition to the ‘cells per mm^2’^ scoring, 7 studies reported only on the overall CD3 positivity/negativity of the IHC staining [40–46], covering a total of 11 different pediatric tumor types (**Fig. 2e**). Overall, most assessed tumor samples were positive for CD3 (mean: 68.44%). Similar to earlier observations, neuroblastoma had a T cell-rich TME with CD3 positivity in all 40 tissue samples analyzed. Diffuse midline glioma and rhabdoid tumors, on the other hand, had a lower CD3 positivity in 79% and 64% respectively. With the ‘percentage of positive cells’ scoring method, we observed a similar trend, where neuroblastoma showed a higher infiltration than brain tumors (**Fig. 2f**). Strikingly, studies reporting T cell densities as cells per field showed an opposite trend [47–49], with medulloblastoma displaying a higher T cell density than osteosarcoma (Supplementary Fig. 1a). These observations were based on single studies and may therefore be less reliable. Taken together, pediatric tumors of the peripheral nerves and soft tissues show a substantial presence of T cells within their TME, whereas brain tumors generally exhibit lower T cell infiltration.

#### 2.2. Distinct NK cell and B cell presence across pediatric tumors

Given the strong association between high B cell and NK cell infiltration and better tumor prognosis [50–52], we characterized the densities of B cells and NK cells within the pediatric TME. Notably, B cell infiltration has been linked to improved response to ICB treatment [53]. However, while T cells were extensively analyzed in the pediatric TME, fewer studies reported on the presence of B cells and NK cells.

For rhabdoid tumors and rhabdomyosarcoma, B cells were detected, however, less frequently than T cells (**Fig. 2g**). Similarly, for neuroblastoma B cells were ∼2.7 times less frequent than T cells (4.6% vs 12.6% positive cells, respectively; **Fig. 2h**), but the reported B cell infiltration was still higher than in brain tumors (gliomas). Also CNS-pnet and osteosarcoma, assessed as ‘cells per mm^2’^, showed low B cell densities (4 and 30 cells/mm^2^, respectively), about 3-4 times fewer B cells than T cells (Supplementary Fig. 1b).The study by Dumars et al. [47], who used ‘cells per field’ as scoring, did not detect any B cells in their analyzed osteosarcoma cohort, which consisted primarily of post-chemotherapy treated samples (Supplementary Fig. 1c). This suggests that the lower numbers or even absence of B cells in the pediatric TME, might be due to the effects of chemotherapy, as chemotherapy is known to reduce B cells numbers [54].

Comparisons of NK cells between studies were particularly challenging due to variability in NK cell markers used. This may have interfered with comparability of detection and quantification of NK cells, as different markers are expressed at varying levels across the stages of NK cell development [55]. In contrast to the relatively low B cell density, NK cells were more often detected in rhabdomyosarcoma and rhabdoid tumors, with a similar fraction of positive samples as T cells (78% and 68%, respectively) (**Fig. 2i**). Also low- and high grade gliomas showed a relatively high NK cell density, with 21% and 12% positive cells, respectively (**Fig. 2j**). Intriguingly, NK cells in low-grade gliomas were detected at >4-fold higher levels than T cells, suggesting that NK cells may be more attractive effector cells for immunotherapies than T cells in these tumors. NK cells were remarkably scarce in neuroblastoma, with less than 1% cells per field testing positive. This contrasts with the high T cell infiltration observed in neuroblastoma, suggesting a preferential infiltration or expansion of T cells over NK cells. This finding is remarkable given the presumed central role of NK cells in the efficacy of Dinutuximab therapy in neuroblastoma through ADCC [56], suggesting potential limitations to its therapeutic efficacy. Additionally, DSRCT and synovial sarcoma showed almost no NK cell infiltration with less than 1 cell per mm^2^ (Supplementary Fig. 1d). In summary, these results indicate that pediatric tumors generally show low B cell infiltration, while NK cell infiltration is relatively high, particularly in brain tumors, underscoring the potential for NK cell-based therapies in these tumors, except for neuroblastoma and DSRCT.

### 3. T cell subsets within the Pediatric Tumor Microenvironment assessed by IHC

Since we observed significant variations in T cell infiltration across different tumor classifications, we assessed the specific T cell subsets present within the pediatric tumors by examining the presence of cytotoxic T cells (CD8^+^), T helper cells (CD4^+^) and T regulatory cells (Tregs; FOXP3^+^).

#### 3.1. Peripheral nerve tumors contain relative high cytotoxic T cell levels

CD8^+^ cytotoxic T cells showed a median density of 28 cells/mm^2^ (range: 1-267 cells/mm^2^; **Fig. 3a**), accounting for approximately 58% of the total T cell population. CD8^+^ T cell levels significantly correlated with general CD3^+^ T cell levels (R=0.89), but ratios differed per tumor type (**Fig. 3b** Supplementary Fig. 2a). Rhabdomyosarcoma tumors showed the lowest proportion of CD8^+^ T cells, comprising 39% of the CD3^+^ T cell population, while ependymoma exhibited the highest at 90% (**Fig. 3b**). For rhabdomyosarcoma the variability observed between studies was again independent of embryonal or alveolar subtypes (**Fig. 3c**). Neuroblastoma studies showed a more consistent CD8^+^ density, except for the study by Mina et al.[31], which reported a considerably lower presence of CD8+ T cells in the TME (**Fig. 3a and d**). Reduced CD8^+^ T cell presence in MYCN-amplified neuroblastomas was reported by all three studies (**Fig. 3d**). We did not observe significant differences of CD8^+^ T cell densities when we grouped tumor types into larger classes, but the trends were similar to CD3^+^ cells, with higher CD8^+^ infiltration in peripheral nerve tumor, and lower infiltration in brain tumor (**Fig. 3e**). CD8^+^ T cell infiltration was consistently higher in adult tumors than pediatric tumors. This suggests that the pediatric TME may be less infiltrated by cytotoxic T cells, potentially limiting the efficacy of immunotherapies developed for adult tumors.

**Figure 3.**
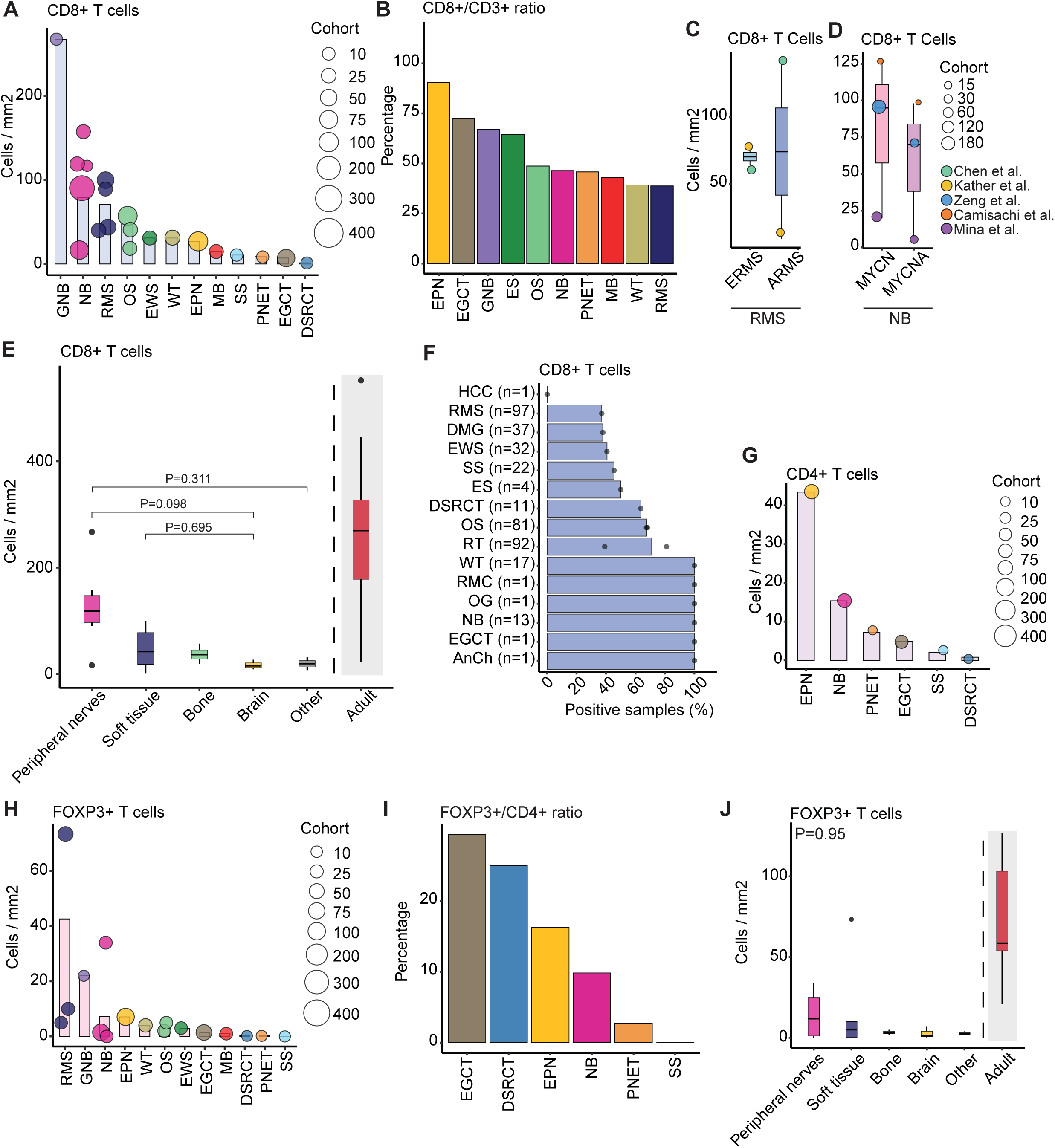
T cell subsets across pediatric brain and solid tumors. (**a**) Bar plot of weighted mean CD8^+^ T cell infiltration per mm^2^ across pediatric (n=12) tumors, with individual (n=12) study distribution. (**b**) CD8/CD3 ratios visualized as percentage CD8^+^ T cells within CD3^+^ T cells across pediatric (n=10) tumors. (**c**) Median CD8^+^ T cell infiltration per mm^2^ in the embryonal (ERMS) and alveolar (ARMS) rhabdomyosarcoma (RMS) subtypes. (**d**) Median CD8^+^ T cell infiltration per mm^2^ in the MYCN non-amplified (MYCN) and MYCN-amplified (MYCNA) neuroblastoma (NB) subtypes. (**e**) Median CD8^+^ T cell infiltration per mm^2^ across tumor categories, based on individual studies. Statistic Kruskal Wallis with Dunn’s and Bonferroni P-value correction was used. (**f**) Percentage of CD8 positive tumor samples across pediatric (n=15) tumors. (**g**) Bar plot of weighted mean CD4^+^ T cell infiltration per mm^2^ across pediatric (n=6) tumors, with individual (n=5) study distribution. (**h**) Bar plot of weighted mean FOXP3^+^ Treg infiltration per mm^2^ across pediatric (n=12) tumors, with individual (n=8) study distribution. (**i**) FOXP3/CD4 ratios visualized as percentage FOXP3^+^ Tregs within CD4^+^ T cells across pediatric (n=6) tumors. (**j**) Median FOXP3^+^ Treg infiltration per mm^2^ across tumor categories, based on individual studies. Statistic Kruskal Wallis with Dunn’s and Bonferroni P-value correction was used. Abbreviations are as follows: MB, medulloblastoma; HGG, high-grade glioma; LGG, low-grade glioma; PNET, primitive neuroectodermal tumors; EPN, ependymoma; RMS, rhabdomyosarcoma; DSRCT, desmoplastic small round cell tumors; NB, neuroblastoma; GNB, ganglioneuroblastoma; OS, osteosarcoma; EWS, Ewing sarcoma; EGCT, extra-cranial germ-cell tumor; WT, Wilms tumor; MM, multiple myeloma; RT, Rhabdoid tumor; AOA, anaplastic oligoastrocytoma; Ch, chordoma; Rb, retinoblastoma; UDS, undifferentiated sarcoma; DMG, diffuse midline glioma; HBL, hepatoblastoma, SS, synovial sarcoma; HCC, hepatocellular carcinoma; RMC, renal medullary carcinoma; OG, oligodendroglioma; AnCh, anaplastic chordoma .

CD8^+^ T cell presence was further supported by the percentage of positive samples analysis, with 6 pediatric tumors, including Wilms tumor and neuroblastoma, showing 100% CD8 positivity in all samples, whereas 9 other pediatric tumors, including brain tumors, exhibited lower positivity (<71% ; **Fig. 3f**). Neuroblastoma also showed higher percentages of CD8^+^ cells as compared to brain tumors, which is in line with the cells per mm^2^ observations (**Supplementary Fig. 2b**). Interestingly, Wilms tumors displayed the highest number of CD8^+^ T cells per field, which contrast with findings based on cells per mm^2^ (**Supplementary Fig. 2c**). This observed discrepancy in CD8^+^ T cell presence may be influenced by chemotherapy-related effects, as the study by Silva et al.[34] (cells/mm^2^) analyzed treatment naïve samples, whereas the study by Mattis et al.[57] (cells/field) comprised both treatment-naïve and post-chemotherapy samples. Altogether, most pediatric tumors show the presence of cytotoxic CD8^+^ T cells within their TME, although their abundance is generally lower compared to adult tumors.

#### 3.2. Consistent T helper and regulatory T cell levels across pediatric tumors

Analysis of CD4^+^ T helper cells revealed a ∼4.7-fold lower density in the pediatric TME compared to CD8^+^ cytotoxic T cells (median 6, range: 1-43 cells/mm^2^ ; **Fig. 3g**). The highest CD4^+^ T cell density was demonstrated in ependymoma (**Fig. 3g**). However, expressed as ‘cells per field’, we observed the opposite with less than 1 CD4^+^ T cell detected per field (**Supplementary Fig. 2d**). Also neuroblastoma showed few CD4^+^ T cells, with only 38.5% of the samples being positive for CD4 (**Supplementary Fig. 2e**).

FOXP3^+^ Tregs, representing 5-10% of the CD4^+^ T helper cells in the circulation [58] are known for their immunosuppressive role within the TME [59,60]. Therefore, we assessed the Treg abundancy to capture the potential immunosuppressive pediatric TME that may hamper immunotherapeutic therapies. Treg densities ranged from 0-43 cells/mm2 (median 3; **Fig. 3h**), with rhabdomyosarcoma showing the highest Treg densities. Of note, this high value was mainly driven by the study of Chen et al. [29], that reported considerably higher densities than the other rhabdomyosarcoma studies. In ependymoma, similar Treg densities were observed as neuroblastoma, but in ependymoma Tregs comprised a higher percentage within the CD4^+^ T helper cells (16% vs 10%, respectively; **Fig. 3i**), suggesting an immunosuppressive TME. No significant difference in Treg densities between tumor categories was observed, but the trend was similar to other T cell analyses, with higher levels in peripheral nerve tumors and lower levels in brain tumors (**Fig. 3j**). We observed a significant positive correlation between Treg and both CD3^+^ and CD8^+^ T cell infiltration, suggesting co-infiltration of Tregs, likely limiting the anti-tumor response of other T cells, including cytotoxic T cells (Supplementary Fig. 2f and 2g).

The presence of Tregs was also found in 57% of rhabdoid tumors and 13% of undifferentiated sarcoma samples (Supplementary Fig. 2h). In terms of ‘percentages of positive cells’ data neuroblastoma showed the highest percentage (2.3%), followed by high- and low-grade glioma (1.6% and 1.1%, respectively ; Supplementary Fig. 2i). In retinoblastoma, 2 Tregs ‘per field’ were found (Supplementary Fig. 2j) Overall, these results indicate that most pediatric tumors are (co-)infiltrated by Tregs, thereby driving a more immunosuppressive TME that suppresses anti-tumor responses and potentially reduces the efficacy of immune therapeutics.

### 4. Tumor-associated macrophages across pediatric solid tumors assessed by IHC

In addition to Tregs, myeloid cells, including macrophages and in the brain microglia, are known key players in shaping the immunosuppressive TME, and their infiltration is often associated with poor prognosis [15,61]. To improve our understanding of the myeloid contribution within the pediatric TME, we examined the presence of CD68^+^ as well as CD163^+^ macrophages/microglia, which we will comprehensively refer to as macrophages.

#### 4.1. Macrophage-dominant tumor microenvironment in all malignant pediatric tumors

Only 4 studies quantified CD68^+^ macrophage infiltration using ‘positive cells per mm^2^’ as scoring method [27,30,32,34], reporting a median density of 218 cells/mm^2^ (range: 114-376 cells/mm^2^; **Fig. 4a)**. The highest CD68^+^ macrophage infiltration was shown in ganglioneuroblastoma, followed by rhabdomyosarcoma and neuroblastoma, with densities similar to those observed in adult tumors (**Fig. 4a** and **Supplementary Fig. 3a**). The CD68^+^ macrophage infiltration significantly correlated with CD3+ T cell infiltration (**Fig. 4b**), suggesting co-infiltration and potential interactions that could negatively influence the immune response. Notably, the median density of CD68^+^ macrophages was found almost 4.5-fold higher than that of CD3^+^ T cells. This macrophage-richness of the pediatric TME was further emphasized by the CD3^+^/CD68^+^ ratios, with ependymoma and rhabdomyosarcoma showing a substantial macrophage predominance, while among pediatric tumors only ganglioneuroblastoma showed a slightly T cell dominance (**Fig. 4c)**. Furthermore, nearly all 7 tumor types analyzed by ‘percentage of positive samples’ showed CD68 positivity (**Fig. 4d**). Osteosarcoma exhibited ∼12-fold higher CD68^+^ macrophage density relative to CD3^+^ T cells when measured by ‘cells per field’ (2 CD3^+^ vs 23 CD68^+^ cells/field; **Supplementary Fig. 1a** and 3b), whereas this difference was only 1.1-fold when assessed by cells per mm^2^. The observed variation in fold change observed may be due to the cohort differences, as the study by Dumars et al. [47] primarily included post-chemotherapy samples, whereas the samples in Silva et al. [34] were treatment-naïve. In neuroblastoma, 6% CD68^+^ macrophages were observed measured as ‘percentage positive cells’ (Supplementary Fig. 3c), which was ∼2-fold higher than the percentage of CD3^+^ positive T cells (11% ; **Fig. 2f**). Altogether, these findings suggests the presence of a myeloid-rich TME in most pediatric tumors that may impact T cell-mediated responses.

**Figure 4.**
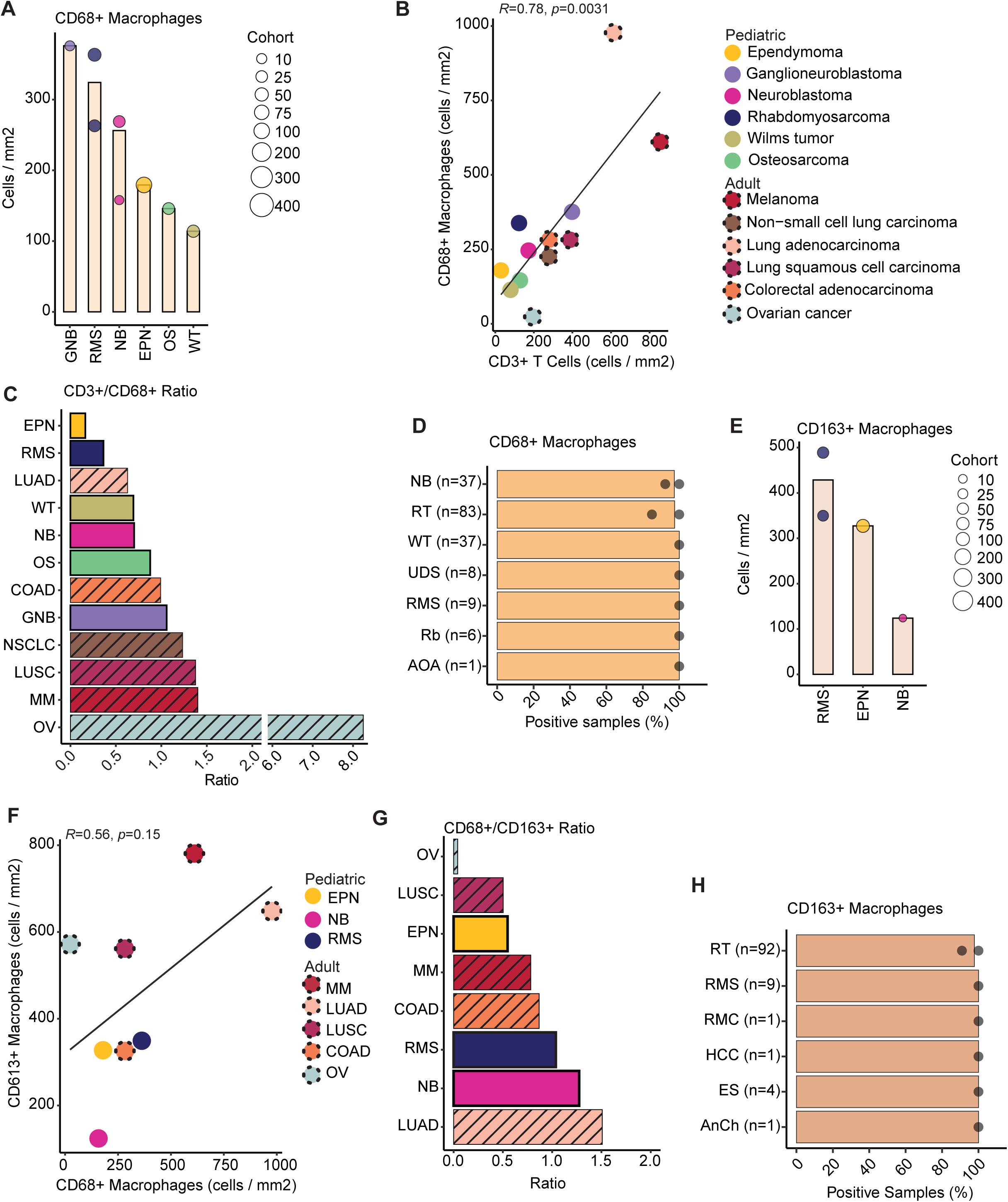
Macrophages infiltration across pediatric brain and solid tumors. (**a**) Bar plot of weighted mean CD68^+^ macrophage infiltration per mm^2^ across pediatric (n=6) tumors, with individual (n=4) study distribution. (**b**) Correlation between CD3^+^ T cells and CD68^+^ macrophages per tumor type. Dashed lines represent adult tumors. (**c**) CD3^+^/CD68^+^ ratios visualized across pediatric (n=6) and adult (n=6) tumors. Diagonal lines represent adult tumors. (**d**) Percentage of CD68 positive tumor samples across pediatric (n=7) tumors. (**e**) Bar plot of weighted mean CD163^+^ macrophage infiltration per mm^2^ across pediatric (n=3) tumors, with individual (n=4) study distribution. (**f**) Correlation between CD68^+^ macrophages and CD163^+^ macrophages per tumor type. Dashed lines represent adult tumors. (**g**) CD68^+^/CD163^+^ ratios visualized across pediatric (n=3) and adult (n=5) tumors. Diagonal lines represent adult tumors. (**h**) Percentage of CD163 positive tumor samples across pediatric (n=6) tumors. Abbreviations are as follows: EPN, ependymoma; RMS, rhabdomyosarcoma; NB, neuroblastoma; GNB, ganglioneuroblastoma; OS, osteosarcoma; WT, Wilms tumor; RT, Rhabdoid tumor; AOA, anaplastic oligoastrocytoma; Rb, retinoblastoma; UDS, undifferentiated sarcoma; SS, synovial sarcoma; HCC, hepatocellular carcinoma; RMC, renal medullary carcinoma; ES, epithelioid sarcoma; AnCh, anaplastic chordoma; MM, melanoma; OV, ovarian cancer; LUSC, lung squamous cell carcinoma; LUAD, lung adenocarcinoma; COAD, colorectal adenocarcinoma; NSCLC, non-small cell lung carcinoma.

#### 4.2. CD163^+^ Macrophage-rich tumor microenvironment in brain and soft tissue tumors

Among the diverse macrophage polarization states, the CD163^+^ macrophages, often characterized as M2-like, are particularly recognized for their potential immunosuppressive phenotype and contribution to metastasis [62]. Rhabdomyosarcoma showed the highest CD163^+^ macrophage infiltration, followed by ependymoma and neuroblastoma, with densities ranging from 124 to 429 cells/mm^2^ (median: 327 cells/mm^2^) (**Fig. 4e**). CD163^+^ macrophage densities were lower than in adult tumors, with neuroblastoma exhibiting the lowest CD163^+^ macrophage density (**Supplementary Fig. 3d**). CD68^+^ and CD163^+^ macrophage densities showed a slightly positive trend, indicating a connection between these subsets, but with substantial difference in proportions between tumors (**Fig. 4f and g**). While neuroblastoma and rhabdomyosarcoma had slightly more CD68^+^ than CD163^+^ macrophages, ependymoma exhibited a greater proportion of CD163^+^ macrophages than CD68^+^ within the TME (**Fig. 4g**). The latter suggests that not all macrophages express CD68, potentially accounting for their underrepresentation in IHC analysis [63]. Similar to CD68^+^ macrophages, CD163^+^ positive macrophages were detected in almost all tumor tissues assessed (**Fig. 4h**). All brain tumors exhibited CD163^+^ macrophages within their TME, as determined by percentage of positive cells, with medulloblastoma showing the lowest density (**Supplementary Fig. 3e)**. Quantified as ‘cells per field’, rhabdoid tumors displayed a CD163^+^ macrophage-rich TME with ∼101 cells per field, whereas osteosarcoma showed a minimal presence of CD163^+^ macrophages with less than 1 cell per field (**Supplementary Fig. 3f**). Altogether, this suggests that, particularly in brain tumors, CD163⁺ macrophages are highly prevalent, potentially contributing to a highly immunosuppressive tumor microenvironment.

### 5. Interactive immune atlas of immunohistochemistry data in R2

To make our concatenated data publicly available to the community, we combined the IHC data (cells/per mm^2^) into an interactive heatmap in R2, which is a publicly accessible online platform (https://hgserver1.amc.nl/cgi-bin/r2/main.cgi?option=imi2_targetmap_v1; map Immune_landscape_mm2(by patient)_v4). This interactive heatmap allows users to view the weighted mean for each cell (sub)type, and for each pediatric brain and solid tumor (sub)type, along with the summary of the studies from which the data was derived (**Fig. 5a and b**). The heatmap includes cohort details for each study – such as cohort size, age range, treatment regimens, and tissue condition – along with an experimental overview and the overall quality score of each study. Additionally, for each study direct links to PubMed are provided for further reference. This interactive heatmaps in R2 provide a comprehensive view of the immune landscape across various pediatric solid tumors as foundation to refine immunotherapeutic strategies for pediatric patients.

**Figure 5.**
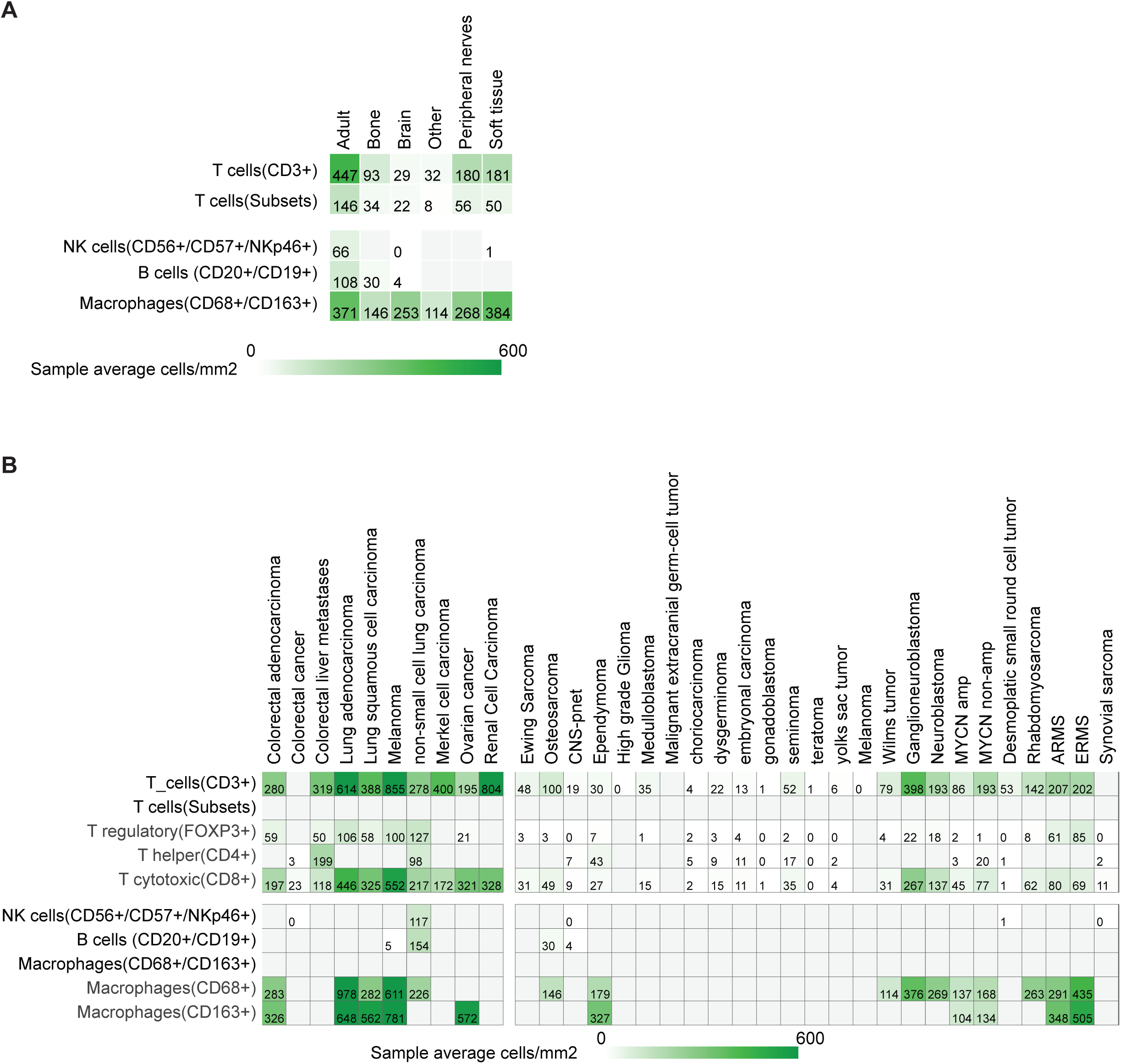
Overview of the pediatric immune landscape in R2. (**a-b**) Class (**a**) and extensive (**b**) overview of the immune cell infiltration per tumor type. Each box represents the weighted mean of the immune cell count per mm^2^, and the color represent the range. The interactive heatmaps are accessible via R2 [https://hgserver1.amc.nl/cgi-bin/r2/main.cgi?option=imi2_targetmap_v1; map Immune_landscape_mm2(by patient)_v4]

### 6. Immune landscape as characterized by single-cell- and single-nucleus RNA-sequencing

In total, 20 studies reported scSeq data [64–83] and 8 studies reported snSeq data [64,80,84–89], collectively examining 1.066.932 cells derived from 272 samples across 16 unique pediatric tumors types (**Fig. 6a**). Consistent with previous studies comparing scSeq and snSeq [39,64,90], we observed that scSeq captures a larger proportion of immune cells (27%) and lower proportion of tumor cells (66%) compared to snSeq (7% and 87%, respectively). Despite the limited studies and relatively small sample sizes available for snSeq, we observed the highest immune proportion for neuroblastoma (13.16%) followed by hepatoblastoma (7.61%) and rhabdomyosarcoma (3.51%) (**Fig. 6b**). This is in line with the IHC data, where we found high T cell and macrophage infiltration for neuroblastoma and rhabdomyosarcoma (**Fig.2b and 4b**). Of note, the limited number of assessed rhabdoid and ETMR samples consisted entirely of tumor cells, although they were not specifically enriched for tumor cells.

**Figure 6.**
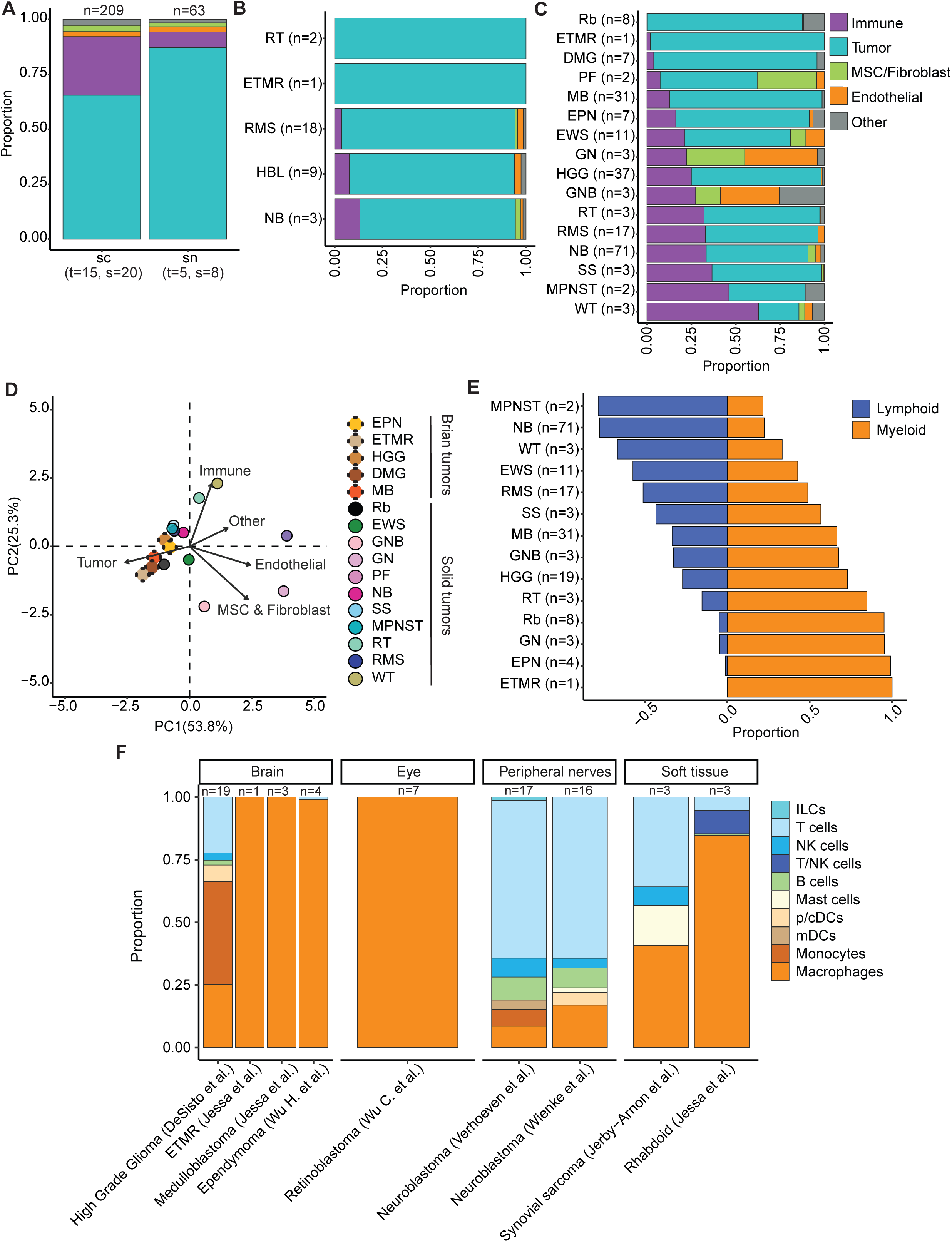
Immune landscape characterization by single-cell and Single-nucleus RNA sequencing. (**a**) Proportion of different cell types per RNA sequencing technique. (**b**) Proportion of different cell types as measured by single-nucleus RNA sequencing per pediatric (n=5) tumors. (**c**) Proportion of different cell types as measured by single-cell RNA sequencing per pediatric (n=16) tumors. (**d**) Principal-component analysis (PCA) plot of pediatric brain (n=5) and solid (n=11) tumors. (**e**) Proportion of myeloid and lymphoid immune cells as measured by single-cell sequencing per pediatric (n=14) tumors, displayed on a divergent scale. (**f**) Proportion of different immune cell types within the immune proportion as measured by single-cell sequencing per pediatric (n=7) tumors. Abbreviations are as follows: Rb, retinoblastoma; ETMR, embryonal tumor with multilayered rosettes; DMG, diffuse midline glioma; PF, plexiform neurofibroma; MB, medulloblastoma; EPN, ependymoma; EWS, Ewing sarcoma; GN, ganglioneuroma; HGG, high-grade glioma; GNB, ganglioneuroblastoma; RT, rhabdoid tumor; RMS, rhabdomyosarcoma; NB, neuroblastoma; SS, synovial sarcoma; MPNST, malignant peripheral nerve sheath tumor; WT, Wilms tumor; HBL, hepatoblastoma.

Next to snSeq data, we examined the pediatric TME based on the available scSeq data. Brain tumors clustered together based on their relatively high proportions of tumor cells (range: 73.15% - 97.99%) and low immune proportions (range: 2.01% - 12.79%), consistent with observations by IHC (**Fig. 6c and d**). Retinoblastoma, a tumor of the eye and, like the brain, an immune privileged site, also clustered together with brain tumors, based on its low immune infiltration (0.22%). Interestingly, neuroblastoma and rhabdomyosarcoma, which had similar immune profiles based on IHC data, also had comparable immune proportion based on scRNAseq (33.31% and 32.98%, respectively). For neuroblastoma, studies varied in tumor cell and immune cell proportions (**Supplementary Fig. 4a**), possibly linked to the differences in treatment regimens and MYCN status between the cohorts. For neuroblastoma, the MYCN- amplified tumors showed lower immune cell proportions than the MYCN non-amplified (11% versus 21%), thereby highlighting their unique TME and confirming observations by IHC (**Supplementary Fig. 4b**). The more benign peripheral nerve tumors ganglioneuroblastoma, ganglioneuroma and plexiform neurofibroma clustered separately from the other pediatric tumors. This separation was driven by their relatively lower tumor cell proportions (range: 0.05% - 54.63% ) and a higher representation of other cell types, including fibroblasts, mesenchymal cells, and vascular cells, supporting the benign characteristics of these tumors. Remarkably, non-small cell lung cancer, a TMB-high adult tumor, clustered together with neuroblastoma and soft tissues sarcomas, showing similar immune proportions (**Supplementary Fig. 4c**).

Focusing on immune cell subsets, we analyzed the ratio of myeloid to lymphoid cells across pediatric tumors. While a substantial variation across tumor type was observed, most tumors, especially brain tumors, showed a myeloid dominant immune profile, with ependymoma, medulloblastoma, retinoblastoma, and EMTR composed (almost) entirely of macrophages (**Fig. 6e and f**). A more lymphoid dominant composition was found for Ewing sarcoma, Wilms tumors, neuroblastoma and malignant peripheral nerve sheath tumors (**Fig. 6e**), aligning with the IHC data. Out of the brain tumors only high grade glioma showed lymphoid infiltration, with some B and NK cell presence, similarly to the IHC data. Remarkably, both neuroblastoma scSeq studies, which exhibited highly similar immune subset proportions, showed a higher NK cell proportion than high grade glioma, which contrasts with the IHC data (**Fig. 2k**). This discrepancy may be linked to the downregulation of typical NK surface markers within the TME, complicating detection by IHC, whereas NK cell detection by scRNAseq based on gene expression is not affected [55,91]. Taken together, brain and eye tumors show similar TME patterns with lower proportion of immune cells that consisted mainly of myeloid cells, suggesting a highly immunosuppressive TME that potentially limits the efficacy of immunotherapeutics.

## Discussion

This systematic review provides a comprehensive overview of the immune landscape across 35 different pediatric brain and solid tumors, integrating data from multiple techniques and scoring methods, including IHC-based quantifications, as well as single-cell- and single-nucleus RNA-seq. The distinct immune landscapes of pediatric brain and solid tumors underscores the importance of tailored immunotherapeutic approaches as direct translation of adult immunotherapy strategies may not be effective. Across pediatric tumors, the predominance of immunosuppressive myeloid populations, coupled with a relative scarcity of CD8^+^ cytotoxic T cells, presents challenges for T cell dependent therapies, which may explain their limited efficacy so far in pediatric patients.

In general, pediatric brain and solid tumors exhibited fewer T cells within their TME than most adults cancers. This discrepancy is most likely due to the overall lower TMB of pediatric tumors, which limits the formation of neoantigens that could be recognized by T cells to mitigate a T cell-mediated antitumor response [7,10,92]. For neuroblastoma and rhabdomyosarcoma, we found notable similarities in their immune landscapes, particularly in terms of their relatively high T cell:macrophage ratio, which could reflect similar interactions with their TME making them more susceptible for immune surveillance. Nevertheless, all pediatric tumors showed a scarcity of CD8^+^ cytotoxic T cells, as compared to adult cancers, thereby reflecting their immune evasive TME. This may have contributed to the lower efficacy of ICB therapies in pediatric than adult patients, as the degree of cytotoxic T cell infiltration largely determines the effectivity of ICB [93–95]. However, given the T cell-rich TME in neuroblastoma and rhabdomyosarcoma, these tumors may benefit from a more general T cell-dependent therapeutic approach such as BiTEs, which are designed to engage CD3^+^ T cells and direct them towards the tumor-associated antigen [96]. Notably, BiTEs (anti-CD3 x anti-GD2) have already shown promising results in refractory/recurrent neuroblastoma and osteosarcoma patients [97]. On the other hand, pediatric brain tumors and solid tumors with relatively low T cell infiltration might benefit more from therapeutic strategies that increases T cell infiltration and activation, such as chemokine modulation [98] and radiotherapy to convert the immunologically “cold” TME into a “hot” TME [99].

Across different methodologies we observed that most pediatric tumors exhibited a macrophage-rich TME. In particular brain tumors showed a higher abundance of myeloid cells relative to lymphoid cells as compared to most pediatric solid tumors. The predominance of CD163^+^ macrophages in brain tumors suggests an immunosuppressive TME that may hamper the efficacy of most currently available immunotherapies, including CAR T cell therapies [100,101]. Given the limited T cell infiltration, pediatric brain tumors may benefit more from therapeutic strategies focusing on the myeloid compartment. Potential strategies to mitigate the highly immunosuppressive M2-like pediatric brain TME, include blocking the CSF-1/CSF1R axis to reduce the recruitment [102], and or reprogramming the M2-like macrophages into a pro-inflammatory M1-like state using BRD4 inhibitors [103]. Another emerging strategy involves the use of CAR macrophages (CAR-M), in which macrophages are reprogrammed to recognize tumor antigens by engineering them to express CARs [104]. In addition to myeloid-targeting therapeutic strategies, pediatric brain tumors may potentially benefit from NK cell-based approaches [105], as most brain tumors showed a relatively high NK presence compared to other pediatric tumors.

A unique strength of our study is that we provide a comprehensive overview of the pediatric TME across 35 tumor types through integration of IHC, scRNA-seq, and snRNA-seq data from a large number of individual studies. By comparing methodologies, we identified key immune landscape differences between pediatric brain and solid tumors. Additionally, our publicly available interactive heatmaps on R2 (https://hgserver1.amc.nl/cgi-bin/r2/main.cgi?option=imi2_targetmap_v1; map Immune_landscape_mm2(by patient)_v4) enhance data accessibility and are aimed at facilitating clinical decision making.

Our study has several limitations, primarily driven by the methodological differences between the studies. Data consistency for IHC was limited by variations in scorings methods, experimental protocols, and immune cell labelling, thereby hampering cross-study comparisons. For sc/snSeq studies, the number of immune cells was often not reported, limiting the number of studies available for analysis, and resulting in fewer tumor types being included. Therefore, standardizing methodologies as well as publicly reporting cell numbers will be crucial for refining our understanding of the pediatric brain and solid tumor TME. Taken together, this systematic review highlights both the strengths and limitations of various immune profiling methods and underscores the need for more integrative approaches that combine spatial, transcriptional, and functional data to fully capture the complexity of the pediatric TME.

We envision this in-depth overview of the immune landscape of pediatric tumors to be at the base of future studies and clinical decision-making, to optimize immunotherapeutic strategies for pediatric tumors. Given the unique immune landscape of each tumor type a one-size-fits-all approach, largely based on adult cancer immunotherapies, will likely not suffice. Rather, tailoring immunotherapy development and selection to specific pediatric tumor types may significantly enhance immunotherapy efficacy.

Altogether, these insights into the unique pediatric TME will be critical for developing effective immunotherapeutic strategies tailored to pediatric patients.

## Acknowledgments

We thank the authors of the original studies included in this systematic review for making their data publicly accessible or for sharing it with us.

---

**Supplementary Figure 1.**
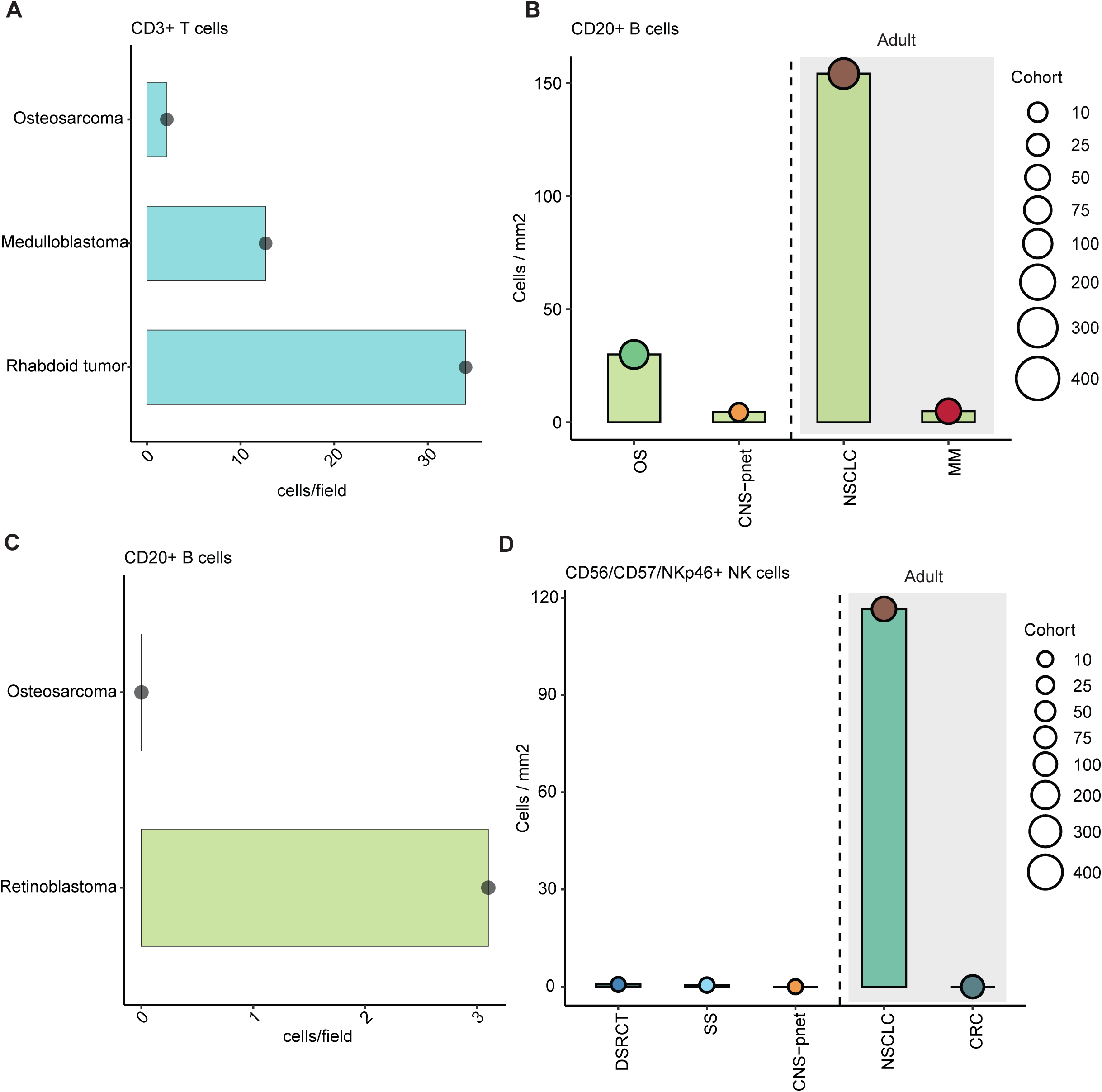
(**a**) CD3^+^ T cells per field across pediatric (n=3) tumors. (**b**) Bar plot of weighted mean CD20^+^ B cell infiltration per mm^2^ across pediatric (n=2) and adult (n=2) tumors. (**c**) CD20^+^ B cells per field across pediatric (n=2) tumors. (**d**) Bar plot of weighted mean CD56^+^/CD57^+^/NKp46^+^ NK cell infiltration per mm^2^ across pediatric (n=3) and adult (n=2) tumors. Abbreviations are as follows: OS, osteosarcoma; CNS-PNET, central nervous system primitive neuroectodermal tumors; SS, synovial sarcoma; DSRCT, desmoplastic small round cell tumors; MM, melanoma; NSCLC; non-small cell lung carcinoma; CRC, colorectal cancer.

**Supplementary Figure 2.**
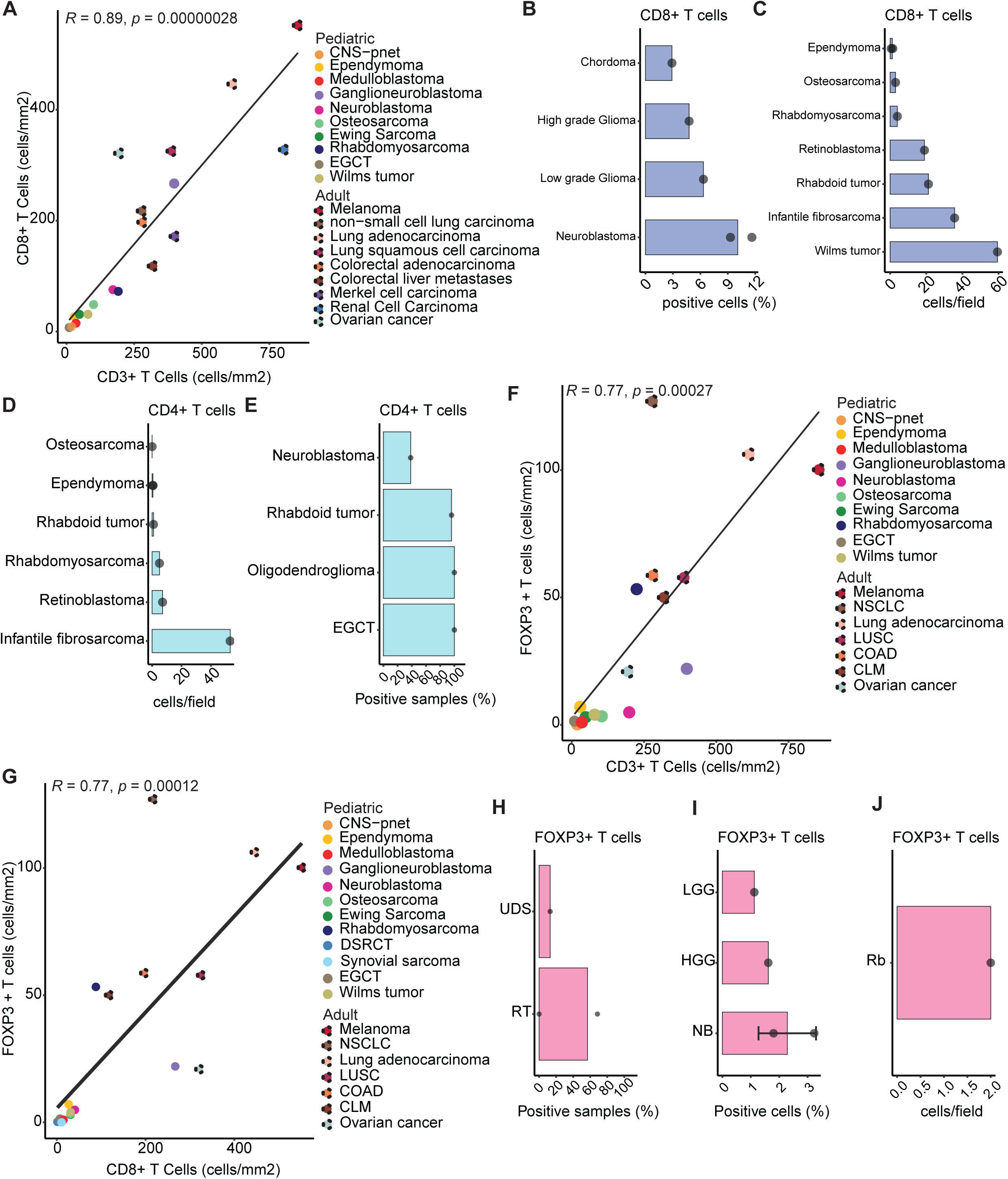
(**a**) Correlation between CD3^+^ T cells and CD8^+^ T cells per tumor type. Dashed lines represent adult tumors. (**b**) Percentage of CD8 positive T cells across pediatric (n=4) tumors. (**c**) CD8^+^ T cells per field across pediatric (n=7) tumors. (**d**) CD4^+^ T cells per field across pediatric (n=6) tumors. (**e**) Percentage of CD4 positive T cells across pediatric (n=4) tumors. (**f**) Correlation between CD3^+^ T cells and FOXP3^+^ Tregs per tumor type. Dashed lines represent adult tumors. (**g**) Correlation between CD8^+^ T cells and FOXP3^+^ Tregs per tumor type. Dashed lines represent adult tumors. (**h**) Percentage of FOXP3 positive tumor samples across pediatric (n=2) tumors. (**i**) Percentage of FOXP3 positive Tregs across pediatric (n=3) tumors. (**j**) FOXP3^+^ Tregs per field in retinoblastoma. Abbreviations are as follows: CNS-PNET, central nervous system primitive neuroectodermal tumors; NB, neuroblastoma; EGCT, extra-cranial germ-cell tumor; UDS, undifferentiated sarcoma; RT, rhabdoid tumors; LGG, low-grade glioma; HGG, high-grade glioma; Rb, retinoblastoma; DSRCT, desmoplastic small round cell tumors; NSCLC; non-small cell lung carcinoma; LUSC, lung squamous cell carcinoma; COAD, colorectal adenocarcinoma; CLM, colorectal liver metastasis.

**Supplementary Figure 3.**
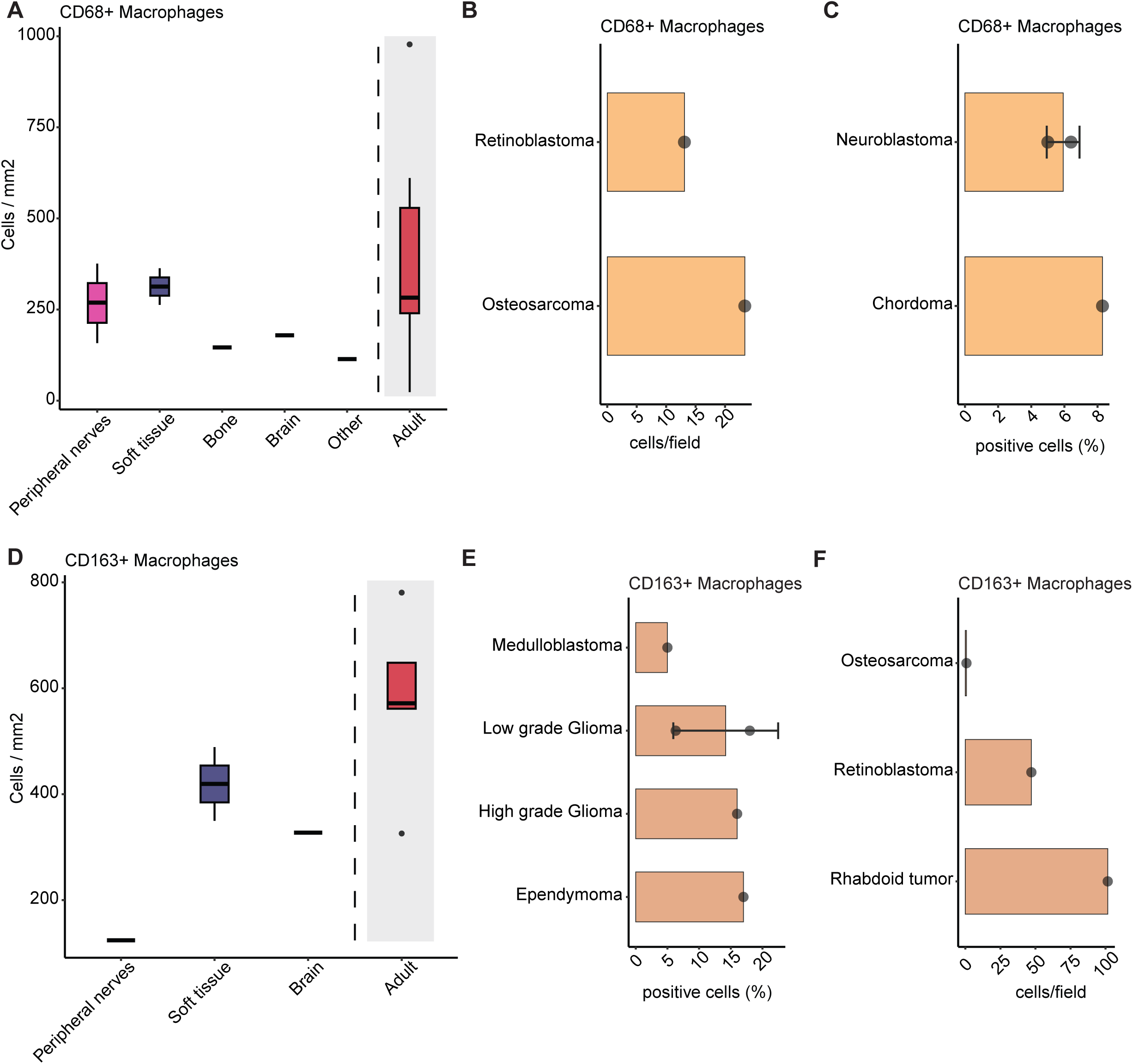
(**a**) Median CD68^+^ macrophage infiltration per mm^2^ across tumor categories, based on individual studies. (**b**) CD68^+^ macrophages per field across pediatric (n=2) tumors. (**c**) Percentage of CD68^+^ positive macrophages across pediatric (n=2) tumors. (**d**) Median CD163^+^ macrophage infiltration per mm^2^ across tumor categories, based on individual studies. (**e**) Percentage of CD163^+^ positive macrophages across pediatric (n=4) tumors. (**f**) CD163^+^ macrophages per field across pediatric (n=3) tumors.

**Supplementary Figure 4.**
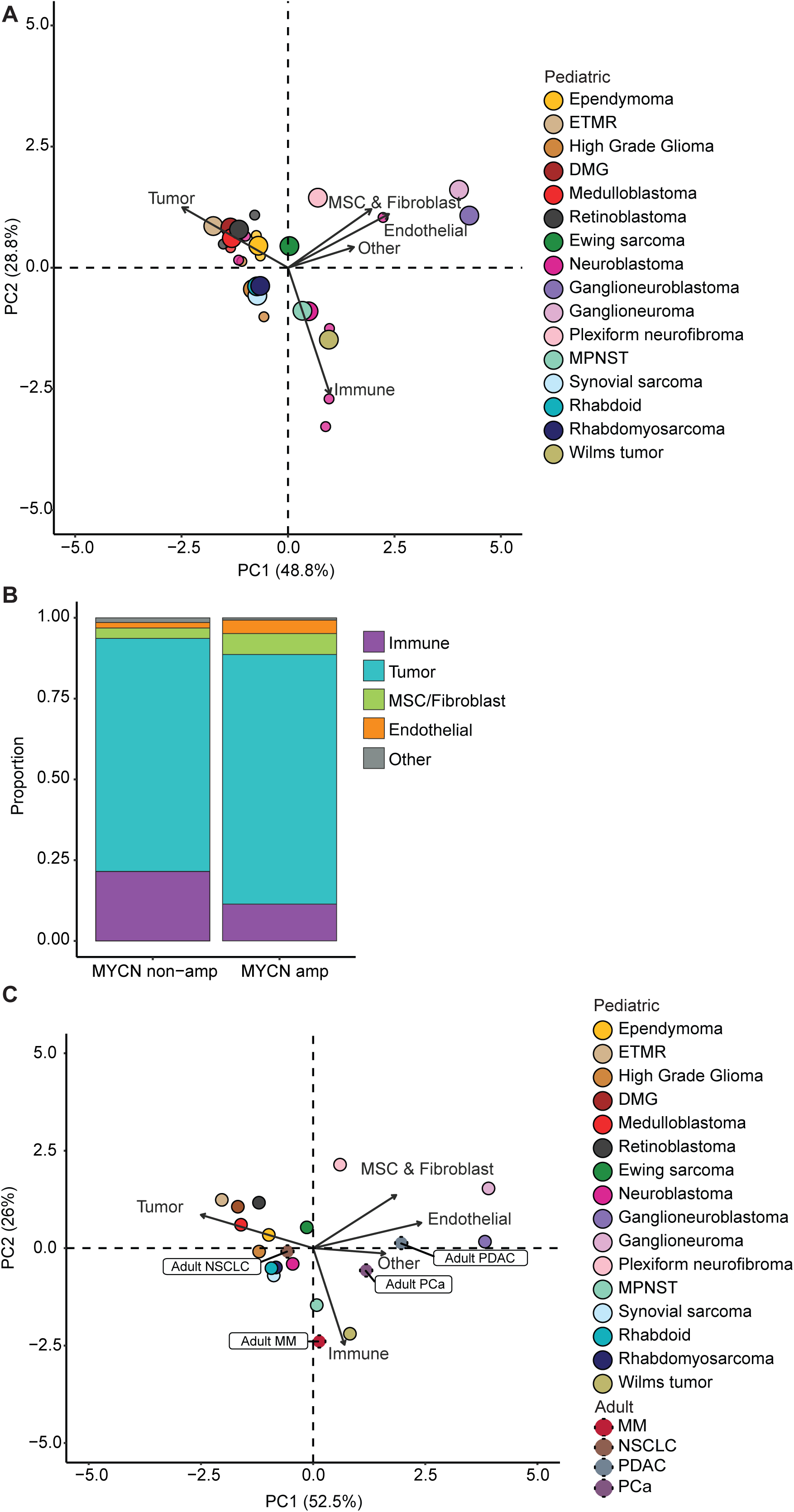
(**a**) Principal-component analysis (PCA) plot of pediatric brain (n=5) and solid (n=11) tumors per study. (**b**) Proportion of different cell types as measured by single-cell RNA sequencing per MYCN non-amplified (MYCN) and MYCN-amplified (MYCNA) neuroblastoma (NB) subtypes (**c**) Principal-component analysis (PCA) plot of pediatric brain (n=5), solid (n=11), and adult (n=4) tumors. Abbreviations are as follows: Rb, retinoblastoma; ETMR, embryonal tumor with multilayered rosettes; DMG, diffuse midline glioma; PF, plexiform neurofibroma; MB, medulloblastoma; EPN, ependymoma; EWS, Ewing sarcoma; GN, ganglioneuroma; HGG, high-grade glioma; GNB, ganglioneuroblastoma; RT, rhabdoid tumor; RMS, rhabdomyosarcoma; NB, neuroblastoma; SS, synovial sarcoma; MPNST, malignant peripheral nerve sheath tumor; WT, Wilms tumor; MM, multiple myeloma; NSCLC, non-small cell lung cancer; PDAC, pancreatic ductal adenocarcinoma; PCa, prostate cancer.

## References

[1] Siegel RL, Giaquinto AN, Jemal A. Cancer statistics, 2024. CA Cancer J Clin 2024;74:12–49. 10.3322/caac.21820.

2. Childhood Cancer Incidence | CureSearch 2014. https://curesearch.org/understanding-childrens-cancer/childhood-cancer-statistics/number-of-diagnoses/ (accessed March 31, 2025).

[3] Zahnreich S, Schmidberger H. Childhood Cancer: Occurrence, Treatment and Risk of Second Primary Malignancies. Cancers 2021;13:2607. 10.3390/cancers13112607.

[4] Pokharkar A, Yadav P, Kandpal DK, Mahajan A, Chowdhary SK. Perioperative and oncologic outcomes of robotic surgery for pediatric solid abdominal tumors: a single-center 10-year experience. Front Pediatr 2025;13. 10.3389/fped.2025.1453718.

[5] Perkins SM, Shinohara ET, DeWees T, Frangoul H. Outcome for Children with Metastatic Solid Tumors over the Last Four Decades. PLoS ONE 2014;9:e100396. 10.1371/journal.pone.0100396.

[6] Landier W, Skinner R, Wallace WH, Hjorth L, Mulder RL, Wong FL, et al. Surveillance for Late Effects in Childhood Cancer Survivors. J Clin Oncol 2018;36:2216–22. 10.1200/JCO.2017.77.0180.

[7] Belgiovine C, Mebelli K, Raffaele A, De Cicco M, Rotella J, Pedrazzoli P, et al. Pediatric Solid Cancers: Dissecting the Tumor Microenvironment to Improve the Results of Clinical Immunotherapy. Int J Mol Sci 2024;25:3225. 10.3390/ijms25063225.

[8] Trubicka J, Grajkowska W, Dembowska-Bagińska B. Molecular Markers of Pediatric Solid Tumors—Diagnosis, Optimizing Treatments, and Determining Susceptibility: Current State and Future Directions. Cells 2022;11:1238. 10.3390/cells11071238.

9. Immunotherapy for Childhood Cancer. Cancer Res Inst n.d. https://www.cancerresearch.org/cancer-types/childhood-cancer (accessed May 19, 2024).

[10] Chen Q, Zhao B, Tan Z, Hedberg G, Wang J, Gonzalez L, et al. Systems-level immunomonitoring in children with solid tumors to enable precision medicine. Cell 2025;188:1425–1440.e11. 10.1016/j.cell.2024.12.014.

[11] Casey DL, Cheung N-KV. Immunotherapy of Pediatric Solid Tumors: Treatments at a Crossroads, with an Emphasis on Antibodies. Cancer Immunol Res 2020;8:161–6. 10.1158/2326-6066.CIR-19-0692.

[12] Devlin JR, Alonso JA, Ayres CM, Keller GLJ, Bobisse S, Vander Kooi CW, et al. Structural dissimilarity from self drives neoepitope escape from immune tolerance. Nat Chem Biol 2020;16:1269–76. 10.1038/s41589-020-0610-1.

[13] Huehls AM, Coupet TA, Sentman CL. Bispecific T cell engagers for cancer immunotherapy. Immunol Cell Biol 2015;93:290–6. 10.1038/icb.2014.93.

[14] Sasidharan Nair V, Elkord E. Immune checkpoint inhibitors in cancer therapy: a focus on T-regulatory cells. Immunol Cell Biol 2018;96:21–33. 10.1111/imcb.1003.

[15] Koo J, Hayashi M, Verneris MR, Lee-Sherick AB. Targeting Tumor-Associated Macrophages in the Pediatric Sarcoma Tumor Microenvironment. Front Oncol 2020;10:581107. 10.3389/fonc.2020.581107.

[16] Yamaguchi Y, Gibson J, Ou K, Lopez LS, Ng RH, Leggett N, et al. PD-L1 blockade restores CAR T cell activity through IFN-γ-regulation of CD163+ M2 macrophages. J Immunother Cancer 2022;10:e004400. 10.1136/jitc-2021-004400.

[17] Ochoa MC, Minute L, Rodriguez I, Garasa S, Perez-Ruiz E, Inogés S, et al. Antibody-dependent cell cytotoxicity: immunotherapy strategies enhancing effector NK cells. Immunol Cell Biol 2017;95:347–55. 10.1038/icb.2017.6.

[18] Kurdi AT, Glavey SV, Bezman NA, Jhatakia A, Guerriero JL, Manier S, et al. Antibody-Dependent Cellular Phagocytosis by Macrophages is a Novel Mechanism of Action of Elotuzumab. Mol Cancer Ther 2018;17:1454–63. 10.1158/1535-7163.MCT-17-0998.

[19] Moher D, Liberati A, Tetzlaff J, Altman DG, Group TP. Preferred Reporting Items for Systematic Reviews and Meta-Analyses: The PRISMA Statement. PLoS Med 2009;6:e1000097. 10.1371/journal.pmed.1000097.

[20] Nabbi A, Beck P, Delaidelli A, Oldridge DA, Sudhaman S, Zhu K, et al. Transcriptional immunogenomic analysis reveals distinct immunological clusters in paediatric nervous system tumours. Genome Med 2023;15:67. 10.1186/s13073-023-01219-x.

[21] Gasparini P, Fortunato O, De Cecco L, Casanova M, Iannó MF, Carenzo A, et al. Age-Related Alterations in Immune Contexture Are Associated with Aggressiveness in Rhabdomyosarcoma. Cancers 2019;11:1380. 10.3390/cancers11091380.

[22] Abro B, Kaushal M, Chen L, Wu R, Dehner LP, Pfeifer JD, et al. Tumor mutation burden, DNA mismatch repair status and checkpoint immunotherapy markers in primary and relapsed malignant rhabdoid tumors. Pathol Res Pract 2019;215:152395. 10.1016/j.prp.2019.03.023.

[23] Kurdi M, Katib Y, Faizo E, Bahakeem B, Alkhotani A, Alkhayyat S, et al. Association Between CD204-Expressed Tumor-Associated Macrophages and MGMT-Promoter Methylation in the Microenvironment of Grade 4 Astrocytomas. World J Oncol 2022;13:117–25. 10.14740/wjon1473.

[24] Meng J, Chen Y, Lu X, Ge Q, Yang F, Bai S, et al. Macrophages and monocytes mediated activation of oxidative phosphorylation implicated the prognosis and clinical therapeutic strategy of Wilms tumour. Comput Struct Biotechnol J 2022;20:3399–408. 10.1016/j.csbj.2022.06.052.

[25] Zhang L, Zhang B, Dou Z, Wu J, Iranmanesh Y, Jiang B, et al. Immune Checkpoint-Associated Locations of Diffuse Gliomas Comparing Pediatric With Adult Patients Based on Voxel-Wise Analysis. Front Immunol 2021;12:582594. 10.3389/fimmu.2021.582594.

[26] Boldrini R, De Pasquale MD, Melaiu O, Chierici M, Jurman G, Benedetti MC, et al. Tumor-infiltrating T cells and PD-L1 expression in childhood malignant extracranial germ-cell tumors. Oncoimmunology 2019;8:e1542245. 10.1080/2162402X.2018.1542245.

[27] Camisaschi C, Renne SL, Beretta V, Rini F, Spagnuolo RD, Tuccitto A, et al. Immune landscape and in vivo immunogenicity of NY-ESO-1 tumor antigen in advanced neuroblastoma patients. BMC Cancer 2018;18:983. 10.1186/s12885-018-4910-8.

[28] Casanova JM, Almeida J-S, Reith JD, Sousa LM, Fonseca R, Freitas-Tavares P, et al. Tumor-Infiltrating Lymphocytes and Cancer Markers in Osteosarcoma: Influence on Patient Survival. Cancers 2021;13:6075. 10.3390/cancers13236075.

[29] Chen L, Oke T, Siegel N, Cojocaru G, Tam AJ, Blosser RL, et al. The Immunosuppressive Niche of Soft-Tissue Sarcomas is Sustained by Tumor-Associated Macrophages and Characterized by Intratumoral Tertiary Lymphoid Structures. Clin Cancer Res Off J Am Assoc Cancer Res 2020;26:4018–30. 10.1158/1078-0432.CCR-19-3416.

[30] Kather JN, Hörner C, Weis C-A, Aung T, Vokuhl C, Weiss C, et al. CD163+ immune cell infiltrates and presence of CD54+ microvessels are prognostic markers for patients with embryonal rhabdomyosarcoma. Sci Rep 2019;9:9211. 10.1038/s41598-019-45551-y.

[31] Mina M, Boldrini R, Citti A, Romania P, D’Alicandro V, Ioris MD, et al. Tumor-infiltrating T lymphocytes improve clinical outcome of therapy-resistant neuroblastoma. Oncoimmunology 2015;4:e1019981. 10.1080/2162402X.2015.1019981.

[32] Nam SJ, Kim Y-H, Park JE, Ra Y-S, Khang SK, Cho YH, et al. Tumor-infiltrating immune cell subpopulations and programmed death ligand 1 (PD-L1) expression associated with clinicopathological and prognostic parameters in ependymoma. Cancer Immunol Immunother CII 2019;68:305–18. 10.1007/s00262-018-2278-x.

[33] Pasqualini C, Rubino J, Brard C, Cassard L, André N, Rondof W, et al. Phase II and biomarker study of programmed cell death protein 1 inhibitor nivolumab and metronomic cyclophosphamide in paediatric relapsed/refractory solid tumours: Arm G of AcSé-ESMART, a trial of the European Innovative Therapies for Children With Cancer Consortium. Eur J Cancer Oxf Engl 1990 2021;150:53–62. 10.1016/j.ejca.2021.03.032.

[34] Silva MA, Triltsch N, Leis S, Kanchev I, Tan TH, Van Peel B, et al. Biomarker recommendation for PD-1/PD-L1 immunotherapy development in pediatric cancer based on digital image analysis of PD-L1 and immune cells. J Pathol Clin Res 2020;6:124–37. 10.1002/cjp2.152.

[35] Thakur MD, Franz CJ, Brennan L, Brouwer-Visser J, Tam R, Korski K, et al. Immune contexture of paediatric cancers. Eur J Cancer Oxf Engl 1990 2022;170:179–93. 10.1016/j.ejca.2022.03.012.

[36] Vermeulen JF, van Hecke W, Spliet WGM, Villacorta Hidalgo J, Fisch P, Broekhuizen R, et al. Pediatric Primitive Neuroectodermal Tumors of the Central Nervous System Differentially Express Granzyme Inhibitors. PLoS ONE 2016;11:e0151465. 10.1371/journal.pone.0151465.

[37] Zeng L, Li S-H, Xu S-Y, Chen K, Qin L-J, Liu X-Y, et al. Clinical Significance of a CD3/CD8-Based Immunoscore in Neuroblastoma Patients Using Digital Pathology. Front Immunol 2022;13:878457. 10.3389/fimmu.2022.878457.

[38] Masih KE, Wei JS, Milewski D, Khan J. Exploring and Targeting the Tumor Immune Microenvironment of Neuroblastoma. J Cell Immunol 2021;3:305–16. 10.33696/immunology.3.111.

[39] Patel AG, Ashenberg O, Collins NB, Segerstolpe Å, Jiang S, Slyper M, et al. A spatial cell atlas of neuroblastoma reveals developmental, epigenetic and spatial axis of tumor heterogeneity. BioRxiv 2024:2024.01.07.574538. 10.1101/2024.01.07.574538.

[40] Apps JR, Hasan F, Campus O, Behjati S, Jacques TS, J Sebire N, et al. The immune environment of paediatric solid malignancies: evidence from an immunohistochemical study of clinical cases. Fetal Pediatr Pathol 2013;32:298–307. 10.3109/15513815.2012.754527.

[41] Bertolini G, Bergamaschi L, Ferrari A, Renne SL, Collini P, Gardelli C, et al. PD-L1 assessment in pediatric rhabdomyosarcoma: a pilot study. BMC Cancer 2018;18:652. 10.1186/s12885-018-4554-8.

[42] Jha P, Manjunath N, Singh J, Mani K, Garg A, Kaur K, et al. Analysis of PD-L1 expression and T cell infiltration in different molecular subgroups of diffuse midline gliomas. Neuropathol Off J Jpn Soc Neuropathol 2019;39:413–24. 10.1111/neup.12594.

[43] Lasky JL, Panosyan EH, Plant A, Davidson T, Yong WH, Prins RM, et al. Autologous Tumor Lysate-pulsed Dendritic Cell Immunotherapy for Pediatric Patients with Newly Diagnosed or Recurrent High-grade Gliomas. Anticancer Res 2013;33:2047–56.

[44] Mochizuki K, Kawana S, Yamada S, Muramatsu M, Sano H, Kobayashi S, et al. Various checkpoint molecules, and tumor-infiltrating lymphocytes in common pediatric solid tumors: Possibilities for novel immunotherapy. Pediatr Hematol Oncol 2019;36:17–27. 10.1080/08880018.2019.1578843.

[45] Trieb K, Lechleitner T, Lang S, Windhager R, Kotz R, Dirnhofer S. Evaluation of HLA-DR expression and T-lymphocyte infiltration in osteosarcoma. Pathol Res Pract 1998;194:679–84. 10.1016/S0344-0338(98)80126-X.

[46] Ildefonso CJ, Kong L, Leen A, Chai SJ, Petrochelli V, Chintagumpala M, et al. Absence of systemic immune response to adenovectors after intraocular administration to children with retinoblastoma. Mol Ther J Am Soc Gene Ther 2010;18:1885–90. 10.1038/mt.2010.139.

[47] Dumars C, Ngyuen J-M, Gaultier A, Lanel R, Corradini N, Gouin F, et al. Dysregulation of macrophage polarization is associated with the metastatic process in osteosarcoma. Oncotarget 2016;7:78343–54. 10.18632/oncotarget.13055.

[48] Diao S, Gu C, Zhang H, Yu C. Immune cell infiltration and cytokine secretion analysis reveal a non-inflammatory microenvironment of medulloblastoma. Oncol Lett 2020;20:397. 10.3892/ol.2020.12260.

[49] Panwalkar P, Pratt D, Chung C, Dang D, Le P, Martinez D, et al. SWI/SNF complex heterogeneity is related to polyphenotypic differentiation, prognosis, and immune response in rhabdoid tumors. Neuro-Oncol 2020;22:785–96. 10.1093/neuonc/noaa004.

[50] Li G-Q, Wang Y-K, Zhou H, Jin L-G, Wang C-Y, Albahde M, et al. Application of Immune Infiltration Signature and Machine Learning Model in the Differential Diagnosis and Prognosis of Bone-Related Malignancies. Front Cell Dev Biol 2021;9:630355. 10.3389/fcell.2021.630355.

[51] Schaafsma E, Jiang C, Cheng C. B cell infiltration is highly associated with prognosis and an immune-infiltrated tumor microenvironment in neuroblastoma. J Cancer Metastasis Treat 2021;7:10.20517/2394-4722.2021.72. 10.20517/2394-4722.2021.72.

[52] Nersesian S, Schwartz SL, Grantham SR, MacLean LK, Lee SN, Pugh-Toole M, et al. NK cell infiltration is associated with improved overall survival in solid cancers: A systematic review and meta-analysis. Transl Oncol 2021;14:100930. 10.1016/j.tranon.2020.100930.

[53] Helmink BA, Reddy SM, Gao J, Zhang S, Basar R, Thakur R, et al. B cells and tertiary lymphoid structures promote immunotherapy response. Nature 2020;577:549–55. 10.1038/s41586-019-1922-8.

[54] Waidhauser J, Schuh A, Trepel M, Schmälter A-K, Rank A. Chemotherapy markedly reduces B cells but not T cells and NK cells in patients with cancer. Cancer Immunol Immunother CII 2020;69:147–57. 10.1007/s00262-019-02449-y.

[55] Abel AM, Yang C, Thakar MS, Malarkannan S. Natural Killer Cells: Development, Maturation, and Clinical Utilization. Front Immunol 2018;9:1869. 10.3389/fimmu.2018.01869.

[56] Mohd AB, Mohd OB, Alabdallat YJ, Al Dwairy SY, Ghannam RA, Hanaqtah BM, et al. Safety and efficacy of dinutuximab in the treatment of neuroblastoma: A review. J Res Med Sci Off J Isfahan Univ Med Sci 2023;28:71. 10.4103/jrms.jrms_727_22.

[57] Mattis AJ, Chen J-F, Gonzalez IA, Rais R, Dehner LP, Pfeifer J, et al. Immune checkpoint markers and tumour mutation burden in Wilms tumour: a study of 59 cases. Pathology (Phila) 2024;56:814–25. 10.1016/j.pathol.2024.03.005.

[58] Lourenço EV, La Cava A. Natural regulatory T cells in autoimmunity. Autoimmunity 2011;44:33–42. 10.3109/08916931003782155.

[59] Brinkrolf P, Landmeier S, Altvater B, Chen C, Pscherer S, Rosemann A, et al. A high proportion of bone marrow T cells with regulatory phenotype (CD4+CD25hiFoxP3+) in Ewing sarcoma patients is associated with metastatic disease. Int J Cancer 2009;125:879–86. 10.1002/ijc.24461.

[60] Maturu P, Jones D, Ruteshouser EC, Hu Q, Reynolds JM, Hicks J, et al. Role of Cyclooxygenase-2 Pathway in Creating an Immunosuppressive Microenvironment and in Initiation and Progression of Wilms’ Tumor. Neoplasia 2017;19:237–49. 10.1016/j.neo.2016.07.009.

[61] Fujiwara T, Fukushi J, Yamamoto S, Matsumoto Y, Setsu N, Oda Y, et al. Macrophage Infiltration Predicts a Poor Prognosis for Human Ewing Sarcoma. Am J Pathol 2011;179:1157–70. 10.1016/j.ajpath.2011.05.034.

[62] Holterhus M, Altvater B, Kailayangiri S, Rossig C. The Cellular Tumor Immune Microenvironment of Childhood Solid Cancers: Informing More Effective Immunotherapies. Cancers 2022;14:2177. 10.3390/cancers14092177.

[63] Tremble LF, McCabe M, Walker SP, McCarthy S, Tynan RF, Beecher S, et al. Differential association of CD68+ and CD163+ macrophages with macrophage enzymes, whole tumour gene expression and overall survival in advanced melanoma. Br J Cancer 2020;123:1553–61. 10.1038/s41416-020-01037-7.

[64] Bonine N, Zanzani V, Van Hemelryk A, Vanneste B, Zwicker C, Thoné T, et al. NBAtlas: A harmonized single-cell transcriptomic reference atlas of human neuroblastoma tumors. Cell Rep 2024;43:114804. 10.1016/j.celrep.2024.114804.

[65] Cuadrado-Vilanova M, Liu J, Paco S, Aschero R, Burgueño V, Sirab N, et al. Identification of immunosuppressive factors in retinoblastoma cell secretomes and aqueous humor from patients. J Pathol 2022;257:327–39. 10.1002/path.5893.

[66] DeMartino J, Meister MT, Visser LL, Brok M, Groot Koerkamp MJA, Wezenaar AKL, et al. Single-cell transcriptomics reveals immune suppression and cell states predictive of patient outcomes in rhabdomyosarcoma. Nat Commun 2023;14:3074. 10.1038/s41467-023-38886-8.

[67] DeSisto J, Donson AM, Griesinger AM, Fu R, Riemondy K, Mulcahy Levy J, et al. Tumor and immune cell types interact to produce heterogeneous phenotypes of pediatric high grade glioma. Neuro-Oncol 2023:noad207. 10.1093/neuonc/noad207.

[68] Dong R, Yang R, Zhan Y, Lai H-D, Ye C-J, Yao X-Y, et al. Single-Cell Characterization of Malignant Phenotypes and Developmental Trajectories of Adrenal Neuroblastoma. Cancer Cell 2020;38:716–733.e6. 10.1016/j.ccell.2020.08.014.

[69] Filbin MG, Tirosh I, Hovestadt V, Shaw ML, Escalante LE, Mathewson ND, et al. Developmental and oncogenic programs in H3K27M gliomas dissected by single-cell RNA-seq. Science 2018;360:331–5. 10.1126/science.aao4750.

[70] Jerby-Arnon L, Neftel C, Shore ME, Weisman HR, Mathewson ND, McBride MJ, et al. Opposing immune and genetic mechanisms shape oncogenic programs in synovial sarcoma. Nat Med 2021;27:289–300. 10.1038/s41591-020-01212-6.

[71] Jessa S, Mohammadnia A, Harutyunyan AS, Hulswit M, Varadharajan S, Lakkis H, et al. K27M in canonical and noncanonical H3 variants occurs in distinct oligodendroglial cell lineages in brain midline gliomas. Nat Genet 2022;54:1865–80. 10.1038/s41588-022-01205-w.

[72] Liu Q, Wang Z, Jiang Y, Shao F, Ma Y, Zhu M, et al. Single-cell landscape analysis reveals distinct regression trajectories and novel prognostic biomarkers in primary neuroblastoma. Genes Dis 2022;9:1624–38. 10.1016/j.gendis.2021.12.020.

[73] Mei S, Alchahin AM, Embaie BT, Gavriliuc IM, Verhoeven BM, Zhao T, et al. Single-cell analyses of metastatic bone marrow in human neuroblastoma reveals microenvironmental remodeling and metastatic signature. JCI Insight 2024:e173337. 10.1172/jci.insight.173337.

[74] Riemondy KA, Venkataraman S, Willard N, Nellan A, Sanford B, Griesinger AM, et al. Neoplastic and immune single-cell transcriptomics define subgroup-specific intra-tumoral heterogeneity of childhood medulloblastoma. Neuro-Oncol 2022;24:273–86. 10.1093/neuonc/noab135.

[75] Verhoeven BM, Mei S, Olsen TK, Gustafsson K, Valind A, Lindström A, et al. The immune cell atlas of human neuroblastoma. Cell Rep Med 2022;3:100657. 10.1016/j.xcrm.2022.100657.

[76] Visser LL, Bleijs M, Margaritis T, van de Wetering M, Holstege FCP, Clevers H. Ewing Sarcoma Single-cell Transcriptome Analysis Reveals Functionally Impaired Antigen-presenting Cells. Cancer Res Commun 2023;3:2158–69. 10.1158/2767-9764.CRC-23-0027.

[77] Wienke J, Visser LL, Kholosy WM, Keller KM, Barisa M, Poon E, et al. Integrative analysis of neuroblastoma by single-cell RNA sequencing identifies the NECTIN2-TIGIT axis as a target for immunotherapy. Cancer Cell 2024;42:283–300.e8. 10.1016/j.ccell.2023.12.008.

[78] Wu C, Yang J, Xiao W, Jiang Z, Chen S, Guo D, et al. Single-cell characterization of malignant phenotypes and microenvironment alteration in retinoblastoma. Cell Death Dis 2022;13:438. 10.1038/s41419-022-04904-8.

[79] Wu H, Fu R, Zhang Y-H, Liu Z, Chen Z-H, Xu J, et al. Single-Cell RNA Sequencing Unravels Upregulation of Immune Cell Crosstalk in Relapsed Pediatric Ependymoma. Front Immunol 2022;13. 10.3389/fimmu.2022.903246.

[80] Jessa S, Blanchet-Cohen A, Krug B, Vladoiu M, Coutelier M, Faury D, et al. Stalled developmental programs at the root of pediatric brain tumors. Nat Genet 2019;51:1702–13. 10.1038/s41588-019-0531-7.

[81] Zhang X, Lou HE, Gopalan V, Liu Z, Jafarah HM, Lei H, et al. Single-cell sequencing reveals activation of core transcription factors in PRC2-deficient malignant peripheral nerve sheath tumor. Cell Rep 2022;40:111363. 10.1016/j.celrep.2022.111363.

[82] Mazuelas H, Magallón-Lorenz M, Fernández-Rodríguez J, Uriarte-Arrazola I, Richaud-Patin Y, Terribas E, et al. Modeling iPSC-derived human neurofibroma-like tumors in mice uncovers the heterogeneity of Schwann cells within plexiform neurofibromas. Cell Rep 2022;38:110385. 10.1016/j.celrep.2022.110385.

[83] Young MD, Mitchell TJ, Vieira Braga FA, Tran MGB, Stewart BJ, Ferdinand JR, et al. Single-cell transcriptomes from human kidneys reveal the cellular identity of renal tumors. Science 2018;361:594–9. 10.1126/science.aat1699.

[84] Bondoc A, Glaser K, Jin K, Lake C, Cairo S, Geller J, et al. Identification of distinct tumor cell populations and key genetic mechanisms through single cell sequencing in hepatoblastoma. Commun Biol 2021;4:1049. 10.1038/s42003-021-02562-8.

[85] Roehrig A, Hirsch TZ, Pire A, Morcrette G, Gupta B, Marcaillou C, et al. Single-cell multiomics reveals the interplay of clonal evolution and cellular plasticity in hepatoblastoma. Nat Commun 2024;15:3031. 10.1038/s41467-024-47280-x.

[86] Bedoya-Reina OC, Li W, Arceo M, Plescher M, Bullova P, Pui H, et al. Single-nuclei transcriptomes from human adrenal gland reveal distinct cellular identities of low and high-risk neuroblastoma tumors. Nat Commun 2021;12:5309. 10.1038/s41467-021-24870-7.

[87] Jansky S, Sharma AK, Körber V, Quintero A, Toprak UH, Wecht EM, et al. Single-cell transcriptomic analyses provide insights into the developmental origins of neuroblastoma. Nat Genet 2021;53:683–93. 10.1038/s41588-021-00806-1.

[88] Stöber MC, Chamorro González R, Brückner L, Conrad T, Wittstruck N, Szymansky A, et al. Intercellular extrachromosomal DNA copy-number heterogeneity drives neuroblastoma cell state diversity. Cell Rep 2024;43:114711. 10.1016/j.celrep.2024.114711.

[89] Patel AG, Chen X, Huang X, Clay MR, Komarova NL, Krasin MJ, et al. The myogenesis program drives clonal selection and drug resistance in rhabdomyosarcoma. Dev Cell 2022;57:1226–1240.e8. 10.1016/j.devcel.2022.04.003.

[90] Slyper M, Porter CBM, Ashenberg O, Waldman J, Drokhlyansky E, Wakiro I, et al. A single-cell and single-nucleus RNA-Seq toolbox for fresh and frozen human tumors. Nat Med 2020;26:792–802. 10.1038/s41591-020-0844-1.

[91] Rebuffet L, Melsen JE, Escalière B, Basurto-Lozada D, Bhandoola A, Björkström NK, et al. High-dimensional single-cell analysis of human natural killer cell heterogeneity. Nat Immunol 2024;25:1474–88. 10.1038/s41590-024-01883-0.

[92] Chan TA, Yarchoan M, Jaffee E, Swanton C, Quezada SA, Stenzinger A, et al. Development of tumor mutation burden as an immunotherapy biomarker: utility for the oncology clinic. Ann Oncol 2019;30:44–56. 10.1093/annonc/mdy495.

[93] Park JA, Cheung N-KV. Limitations and opportunities for immune checkpoint inhibitors in pediatric malignancies. Cancer Treat Rev 2017;58:22–33. 10.1016/j.ctrv.2017.05.006.

[94] Jiang P, Gu S, Pan D, Fu J, Sahu A, Hu X, et al. Signatures of T cell dysfunction and exclusion predict cancer immunotherapy response. Nat Med 2018;24:1550–8. 10.1038/s41591-018-0136-1.

[95] Blumenthal DT, Yalon M, Vainer GW, Lossos A, Yust S, Tzach L, et al. Pembrolizumab: first experience with recurrent primary central nervous system (CNS) tumors. J Neurooncol 2016;129:453–60. 10.1007/s11060-016-2190-1.

[96] Gupta S, Rau RE, Kairalla JA, Rabin KR, Wang C, Angiolillo AL, et al. Blinatumomab in standard risk pediatric B-acute lymphoblastic leukemia. N Engl J Med 2025;392:875–91. 10.1056/NEJMoa2411680.

[97] Yankelevich M, Thakur A, Modak S, Chu R, Taub J, Martin A, et al. Targeting refractory/recurrent neuroblastoma and osteosarcoma with anti-CD3×anti-GD2 bispecific antibody armed T cells. J Immunother Cancer 2024;12:e008744. 10.1136/jitc-2023-008744.

[98] Lim RJ, Salehi-Rad R, Tran LM, Oh MS, Dumitras C, Crosson WP, et al. CXCL9/10-engineered dendritic cells promote T cell activation and enhance immune checkpoint blockade for lung cancer. Cell Rep Med 2024;5:101479. 10.1016/j.xcrm.2024.101479.

[99] Sevenich L. Turning “Cold” Into “Hot” Tumors—Opportunities and Challenges for Radio-Immunotherapy Against Primary and Metastatic Brain Cancers. Front Oncol 2019;9:163. 10.3389/fonc.2019.00163.

[100] Hadiloo K, Taremi S, Heidari M, Esmaeilzadeh A. The CAR macrophage cells, a novel generation of chimeric antigen-based approach against solid tumors. Biomark Res 2023;11:103. 10.1186/s40364-023-00537-x.

[101] Patterson JD, Henson JC, Breese RO, Bielamowicz KJ, Rodriguez A. CAR T Cell Therapy for Pediatric Brain Tumors. Front Oncol 2020;10:1582. 10.3389/fonc.2020.01582.

[102] Wen J, Wang S, Guo R, Liu D. CSF1R inhibitors are emerging immunotherapeutic drugs for cancer treatment. Eur J Med Chem 2023;245:114884. 10.1016/j.ejmech.2022.114884.

[103] Wang S, Wang J, Chen Z, Luo J, Guo W, Sun L, et al. Targeting M2-like tumor-associated macrophages is a potential therapeutic approach to overcome antitumor drug resistance. Npj Precis Oncol 2024;8:1–19. 10.1038/s41698-024-00522-z.

[104] Pierini S, Gabbasov R, Oliveira-Nunes MC, Qureshi R, Worth A, Huang S, et al. Chimeric antigen receptor macrophages (CAR-M) sensitize HER2+ solid tumors to PD1 blockade in pre-clinical models. Nat Commun 2025;16:706. 10.1038/s41467-024-55770-1.

[105] Fares J, Davis ZB, Rechberger JS, Toll SA, Schwartz JD, Daniels DJ, et al. Advances in NK cell therapy for brain tumors. Npj Precis Oncol 2023;7:1–17. 10.1038/s41698-023-00356-1.

